# Development of opioid-induced hyperalgesia depends on reactive astrocytes controlled by Wnt5a signaling

**DOI:** 10.1101/2021.06.19.449129

**Authors:** Xin Liu, Chilman Bae, Bolong Liu, Yongmei Zhang, Xiangfu Zhou, Donghang Zhang, Cheng Zhou, Adriana DiBua, Livia Schutz, Martin Kaczocha, Michelino Puopolo, Terry P. Yamaguchi, Jin Mo Chung, Shao-Jun Tang

## Abstract

Opioids are the frontline analgesics for managing various types of pain. Paradoxically, repeated use of opioid analgesics may cause an exacerbated pain state known as opioid-induced hyperalgesia (OIH), which significantly contributes to dose escalation and consequently opioid overdose. Neuronal malplasticity in pain circuits has been the predominant proposed mechanism of OIH expression. Although glial cells are known to become reactive in OIH animal models, their biological contribution to OIH remains to be defined and their activation mechanism remains to be elucidated. Here, we show that reactive astrocytes (a.k.a. astrogliosis) are critical for OIH development in both male and female mice. Genetic ablation of astrogliosis inhibited the expression of OIH and morphine-induced neural circuit polarization (NCP) in the spinal dorsal horn (SDH). We found that Wnt5a is a neuron-to-astrocyte signal that is required for morphine-induced astrogliosis. Conditional knock-out of Wnt5a in neurons or its co-receptor ROR2 in astrocytes blocked not only morphine-induced astrogliosis but also OIH and NCP. Furthermore, we showed that the Wnt5a-ROR2 signaling-dependent astrogliosis contributes to OIH via inflammasome-regulated IL-1β. Our results reveal an important role of morphine-induced astrogliosis in OIH pathogenesis and elucidate a neuron-to-astrocyte intercellular Wnt signaling pathway that controls the astrogliosis.

## INTRODUCTION

Opioid analgesics such as morphine are the gold standard for treating severe pain. However, long-term use of opioid analgesics often leads to decrease of their efficacy in pain control. Both opioid tolerance and opioid-induced hyperalgesia (OIH) contribute to the decrease of efficacy and consequently dose escalation and overdose^1^. While development of tolerance is manifested by needed dose increase in order to achieve analgesia^2, 3^, OIH by paradoxical increase in pain by use of opioid analgesics^4–7^.. Opioids tolerance and OIH are closely related and likely share specific molecular pathways for their expression^2 8^. OIH is particularly problematic because further opioid prescribing is largely futile^1^. Despite its clinical significance, the mechanisms of OIH pathogenesis are still poorly understood, hampering the development of effective interventions.

Several forms of malplasticity in pain neuronal circuits are implicated in OIH pathogenesis. These include opioid-induced sensitization of nociceptors, sensitization of second order neurons in the spinal cord, and enhanced descending facilitation^7, 9^. Long-term potential (LTP) of synapses between nociceptive C fibers and second order neurons in the spinal cord dorsal horn is considered as a critical synaptic mechanism facilitating OIH expression^9, 10^. Indeed, LTP and OIH appear to use similar molecular signaling pathways that involve NMDA receptors, CaMKII and BDNF to support their expression^9^. LTP might provide a synaptic support to enhance communication between pronociceptive neurons. It is less clear whether other forms of synaptic plasticity in the pain neuronal circuits contribute to OIH expression.

In addition to neuronal malplasticity, opioids also induce reactions of glia, including microglia and astrocytes^9^. Microglia were reported to play a critical role in OIH ^11^ [but see recent studies by Liu et al.^12^]. Astrocytes are activated in the spinal cords following different paradigms of opioid administration^13, 14^. However, despite of suggestive evidence, the potential role of reactive astrocytes in OIH expression has not been conclusively established, and the mechanism by which opioids cause the activation of astrocytes has not elucidated.

Astrocytes intimately interact with neurons, especially with synapses to form tripartite synaptic structures^15^. By interacting with synapses, astrocytes are positioned to sense and respond to synaptic activity-regulated signals from neurons. Because reactive astroglia are widely observed in pain pathways of various models of pathological pain^16^, it is tempting to conceive that pain-associated hyper-synaptic activity may modulate the activation of astrocytes. However, such neuron-to-astrocyte signals have not been elucidated, and their contribution to OIH expression has not been established.

On the other hand, astrocytes may also regulate neuronal functions. Under normal physiological conditions, astrocytes play essential roles in regulating central nervous system homeostasis^17^, plasticity of neural circuits^18^, and synaptic transmission ^15, 19, 20^. However, how reactive astrocytes may regulate synaptic transmission and plasticity remains elusive. In particular, it is unclear whether reactive astrocytes contribute to malsynaptic plasticity of pain circuits associated with OIH development. Although reactive astrocytes are thought to dysregulate neuronal circuits by secreting bioactive molecules such as chemokines and cytokines, the involvement of such a mechanism in OIH pathogenesis has not been tested directly.

In this study, we use a mouse OIH model to show that reactive astrocytes are critical for morphine-induced OIH and malsynaptic plasticity in the SDH. We also demonstrate that the morphine-elicited astrocyte activation is controlled by a neuron-to-astrocyte intercellular Wnt5a-ROR2 signaling pathway. We further elucidate that the reactive astrocytes promote OIH and malsynaptic plasticity via inflammasome-regulated IL-1β.

## MATERIALS AND METHODS

### Animals

Adult male and female C57BL/6 mice (9–10 weeks of age) and GFAP-TK (B6.Cg-Tg(Gfap-TK)7.1Mvs/J) mice^21^ were obtained from Jackson Laboratory (Bar Harbor, ME). Floxed Wnt5a mice were generated as previously described^22^. To delete Wnt5a in neurons, floxed Wnt5a mice were crossed with B6.Cg-Tg (Syn1-cre) 671Jxm/J mice^23^ (Jackson Laboratory) to generate Syn1-cre/Wnt5a^flox/flox^ mice. To delete Wnt5a in neural stem cells, floxed Wnt5a mice were crossed with B6.Cg-Tg(Nes-cre)1Kln/J mice^24^ (Jackson Laboratory) to generate nestin-cre/Wnt5a^flox/flox^ mice. To delete ROR2 in astrocytes, floxed ROR2 mice^25^ (Jackson Laboratory) were crossed with B6.Cg-Tg(GFAP-cre)77.6Mvs/2J mice^26^ (Jackson Laboratory) to generate GFAP-cre/ROR2^flox/flox^ mice. Animal protocols were approved by the Institutional Animal Care and Use Committee of the University of Texas Medical Branch.

### Materials

Morphine sulfate was purchased from West-Ward (Eatontown, NJ). PLX5622-containing rodent diet (1 kg containing 1200 mg PLX5622) was purchased from Research Diets (New Brunswick, NJ). Ganciclovir sodium was purchased from Advanced ChemBlocks Inc. (Burlingame, CA). Interleukin 1 receptor antagonist (IL-1Ra) protein was purchased from R&D Systems Inc. (Minneapolis, MN). AC-YVAD-CMK, a selective inhibitor of caspase-1, JNK Inhibitor II (SP600125), NMDA receptor antagonist APV, selective astrocytic toxin L-α-aminoadipate (L-AA) as well as Gabapentin were purchased from Sigma-Aldrich (St. Louis, MO). Caspase-1 siRNA was purchased from Thermo Fisher Scientific (Waltham, MA). Antibodies used for immunoblotting were anti-IBa1 (1:1000; Wako, 016-20001); anti-GFAP (1:1000; Millipore, MAB360), anti-β-actin (1:1000; Santa Cruz Biotechnology, sc-1616-R), anti-IL-1β (1:500; Novus Biologicals, NB600-633), anti-Wnt5a (1:1000; R&D, MAB645), anti-caspase-1 (1:1000; Adiogen Corporation, AG-20B-0042). Antibodies used for immunohistochemistry were anti-GFAP (1:200; Millipore, AB5541) and anti-IL-1β (1:200; Novus Biologicals, NB600-633).

### Drug administration

Mice were intraperitoneally (i.p.) injected with morphine sulfate (20 mg/kg) daily for 4 consecutive days to establish morphine-induced hyperalgesia. For microglial ablation, C57BL/6 mice were fed a rodent diet containing PLX5622 starting 5 days prior to morphine administration and continuing until the end of experimentation. To ablate reactive astrocytes, GFAP-TK or control WT mice were administered ganciclovir (5 mg/kg) by intrathecal (i.t.) injection for 2 consecutive days during the inductive or maintenance phase of opioid-induced hyperalgesia. AC-YVAD-CMK (4 nmol/kg) or siRNA (3 nmol/kg) were injected (i.t.) daily for the first 4 days. IL-1Ra (20 μg/kg) was injected (i.t.) daily for the first 3 days. Gabapentin (100mg/kg) was injected (i.p.), following the schedule described in (conditioned place preference) CPP section. APV (i.t., 5μg/5μl), SP600125 (i.t., 10μg/5μl), or L-AA (i.t., 100nmol/5μl) was administered 30 min before morphine injection.

### Measurement of mechanical nociception by von Frey test

Mechanical nociceptive hypersensitivity in mice was measured as previously described^27^. The plantar surface of the hind paw of mice in a resting state was stimulated with calibrated von Frey filaments (Stoelting, Wood Dale, IL), and paw withdrawal threshold was determined using the Dixon up and down paradigm. All tests were conducted 2 h prior to drug administration by an experimenter blind to the treatments received by individual animals.

### Conditioned place preference (CPP)

Conditioned place preference (CPP) tests were performed on WT mice with established OIH by repeated morphine administration (i.p., 20 mg/kg daily for 7 consecutive days), according to described procedures^28^ with slight modifications. The procedure had three phases: preconditioning, conditioning, and testing. During the preconditioning phase, mice were placed into standard place conditioning boxes with two chambers (Med Associates, Fairfax, VT, US) for 30 minutes daily for three consecutive days and the door between two chambers was keeping open for free access. The two chambers are differentiated by wall colors (black or white) and floor panels (metal mesh or bar). The natural place preference of mice was determined on the fourth day by recording the time spent in different chambers in 900 seconds duration. In this study, gabapentin was pairing with non-preferred chamber and saline was pairing with preferred chamber. During conditioning phase, mice were administered either gabapentin (i.p., 100mg/kg) or saline once a day with at least 5 hours injection interval and immediately confined to assigned conditioning chamber with a closed door for 30 minutes after injection. Conditioning took three consecutive days and was followed by testing phase. In testing session, mice were placed into the conditioning box with an opening door in the absence of gabapentin or saline injection. To induce OIH, morphine (20mg/kg) or saline (control) were intraperitoneally administered to mice once daily from the day when preconditioning started to the end of conditioning for total 7 days. Morphine or control saline injection was performed after everyday CPP completion. CPP was measure by the time spent in the drug-paired side during testing phase minus the time spent in the drug-paired side during preconditioning phase.

### Western blotting analysis

Mice were anesthetized with 3% isoflurane and sacrificed to collect L4-L6 lumbar spinal cord segments. Tissue was homogenized in RIPA lysis buffer (1% Nonidet P-40, 50 mM Tris-HCl pH 7.4, 150 mM NaCl, 0.5% sodium deoxycholate, and 1 mM EDTA, pH 8.0) with a protease inhibitor cocktail (Sigma). BCA Protein Assay kits (Thermo Fisher) were used to determine protein concentrations. Equal amounts of protein (2 µg) were loaded and separated by 12% SDS-PAGE and transferred to polyvinylidene fluoride membranes. The membranes were blocked in 5% nonfat milk in Tris-buffered saline with 0.1% Tween 20 (TBST) for 1 h at room temperature and then incubated with primary antibodies in TBST buffer overnight at 4°C. After washing with TBST buffer (three times for 10 min at room temperature), membranes were incubated with HRP-conjugated secondary antibody. Enhanced chemiluminescence kits (Pierce) were used to visualize protein bands, and NIH ImageJ software was used for quantification. β-actin was used as a loading control.

### Immunohistochemistry

Mice were anesthetized with 3% isoflurane and transcardially perfused with 20 ml of 0.01 M phosphate-buffered saline (PBS; 0.14 M NaCl, 0.0027 M KCl, 0.010 M PO_4_^3^) followed by 30 ml of 4% paraformaldehyde (PFA) in 0.01 M PBS. L4 and L5 lumbar spinal cord segments were fixed in 4% PFA solution for 12 h at 4°C, dehydrated with 30% sucrose solution in PBS for 24 h at 4°C, and embedded in optimal cutting temperature medium (Tissue-Tek). Tissue was sectioned (15 μm) on a cryostat (Leica CM 1900) and mounted onto Superfrost Plus microscope slides. Sections were incubated in blocking solution containing 5% bovine serum albumin (BSA) and 0.3% Triton X-100 in 0.01 M PBS for 1 h at room temperature and then in chicken anti-GFAP (1:200; Millipore, AB5541) or rabbit IL-1β (1:200; Novus Biologicals, NB600-633) in blocking solution overnight. Sections were washed with 0.01 M PBS (three times for 10 min at room temperature) and incubated with FITC-or Texas Red-conjugated secondary antibody (1:200; Jackson ImmunoResearch Laboratories) in dilution buffer (1% BSA and 0.3% Triton X-100 in 0.01 M PBS) before mounting. Finally, sections were stained with DAPI (Sigma) to visualize nuclei. Animal species-matched IgG was used as a negative control for primary antibodies. Images were collected on a confocal microscope (model A1, Nikon s).

### Patch-clamp recording of dorsal horn neurons in ex vivo spinal cord slices

Spinal cord slices were prepared as previously described^29^. Briefly, the spinal cord was sliced transversely at a thickness of 350 µm using a vibratome (Leica VT1200S, Buffalo Grove, IL) in cold (4°C) N-methyl-D-glucamine (NMDG) solution (93 mM NMDG, 2.5 mM KCl, 1.2 mM NaH_2_PO_4_, 30 mM NaHCO_3_, 20 mM HEPES, 25 mM glucose, 5 mM sodium ascorbate, 2 mM thiourea, 3 mM sodium pyruvate, 10 mM MgSO_4_, and 0.5 mM CaCl_2_, pH 7.4) saturated with 95% O_2_ and 5% CO_2_. Whole-cell recordings were performed from randomly selected neurons in lamina II in artificial cerebrospinal fluid (ACSF; 124 mM NaCl, 2.5 mM KCl, 1.2 mM NaH_2_PO_4_, 24 mM NaHCO_3_, 5 mM HEPES, 12.5 mM glucose, 2 mM MgSO_4_, and 2 mM CaCl_2_, pH 7.4) using a Multiclamp 700B amplifier, DigiDATA, and pClamp software (version 10.6. Molecular Device, Sunnyvale, CA) with a 10-kHz sampling rate and 2-kHz filtering rate. Patch pipettes (4–8 MΩ) were filled with internal solution (120 mM K-gluconate, 10 mM KCl, 2 mM Mg-ATP, 0.5 mM Na-GTP, 0.5 mM EGTA, 20 mM HEPES, and 10 mM phosphocreatine, pH 7.3). After achieving a whole-cell recording configuration, spontaneous excitatory postsynaptic currents (sEPSCs) were recorded for 60 s at - 65 mV in ACSF. Evoked EPSCs (eEPSCs) were elicited by focal electrical stimulation in the vicinity of recorded neurons with a metal bipolar electrode (MicroProbes, Gaithersburg, MD). Test pulses were delivered for 0.5 ms at 5-s intervals, with stimulation intensities ranging from 20–200 µA (20-µA steps). Recordings were performed only when eEPSCs were monosynaptic, based on characteristic waveforms with short latency, a single peak, and stable responses to repeated stimuli. Recordings showing polysynaptic responses were disregarded. Neurons were characterized by their action potential firing pattern upon depolarizing current injections in current clamp mode^30^.

### snRNA-seq analysis of human spinal lumbar spinal cords

The snRNA-seq results for Wnt5a in neurons and ROR2 in astrocytes came from the deposited large datasets that published in bioRxiv^31, 32^. snRNA-seq was performed as described^31^. Briefly, the lumber spinal cords were acutely isolated from adult brain-dead human donors. The nucleus was isolated using a Nucleus Isolation Kit (catalogue no. 52009-10, SHBIO, China) according to the manufacturer’s protocols. The nuclei suspension was loaded onto a Chromium single cell controller (10x Genomics) to produce single-nucleus gel beads in the emulsion (GEM) using single cell 3’ Library and Gel Bead Kit V3.1 (10x Genomics, 1000075) and Chromium Single Cell B Chip Kit (10x Genomics, 1000074) according to the manufacturer’s instructions. The library was sequenced using a Novaseq 6000 sequencing platform (Illumina) with a depth of at least 30,000 reads per nucleus with a paired-end 150 bp (PE150) reading strategy. Cell clustering was performed using Seurat 3.0 (R package).

### Real-time PCR

RNA was extracted using the PureLink RNA Mini Kit (ThermoFisher) following the manufacturer’s instructions. qPCR was performed using PowerUp SYBR green (ThermoFisher) on a StepOne Plus instrument (Applied Biosystems) and negative control reactions lacking cDNA were included. Quantification was performed using the 2-ΔΔCt method and normalized to β-actin. Primers for IL-1β (NM_008361): Forward GCTTCAGGCAGGCAGTATC; Reverse AGGATGGGCTCTTCTTCAAAG. Primers for Caspase-1 (NM_009807): Forward AGGAATTCTGGAGCTTCAATCAG; Reverse TGGAAATGTGCCATCTTCTTT. Primers for β-actin: Forward GACGGCCAGGTCATCACTAT; Reverse CGGATGTCAACGTCACACTT.

### Statistical analysis

Statistical analysis was conducted with Prism 5 (GraphPad) software. Data are shown as mean ± standard error of the mean (SEM). One-way ANOVA was used for immunoblotting data, and two-way ANOVA with Bonferroni post hoc tests were used for pain behavior data. Electrophysiological data are expressed as mean ± SEM with n indicating the number of cells and N indicating the number of animals. sEPSC frequency and eEPSC amplitude were analyzed offline using Clampfit software (version 11, Molecular Devices, CA). sEPSC events were detected using the template event detection method. Electrophysiological data were analyzed using two-way ANOVA followed by Holm-Sidak multiple comparison tests. For all tests, p < 0.05 was considered statistically significant. Origin (version PRO 2020, OriginLab, Northampton, MA) was used to analyze data.

## RESULTS

### Reactive astrocytes play essential role in OIH expression and maintenance

To induce OIH, we repeatedly injected wild-type (WT; C57BL) mice with morphine (i.p.; 20 mg/kg/day for 4 days). We observed the expression of mechanical OIH following the morphine administration (Fig. 1C), as reported previously ^5, 33^. To test if morphine-treated mice develop spontaneous pain, we performed conditioned place preference (CPP) tests with gabapentin, an analgesic suggested to treat OIH in patients^34^. The result showed that the morphine-treated mice spent significantly longer time in the chamber in which gabapentin was administered (i.p., 100mg/kg). These data suggest the development of spontaneous pain in morphine-treated mice (Supplemental Fig 1).

**Figure 1.**
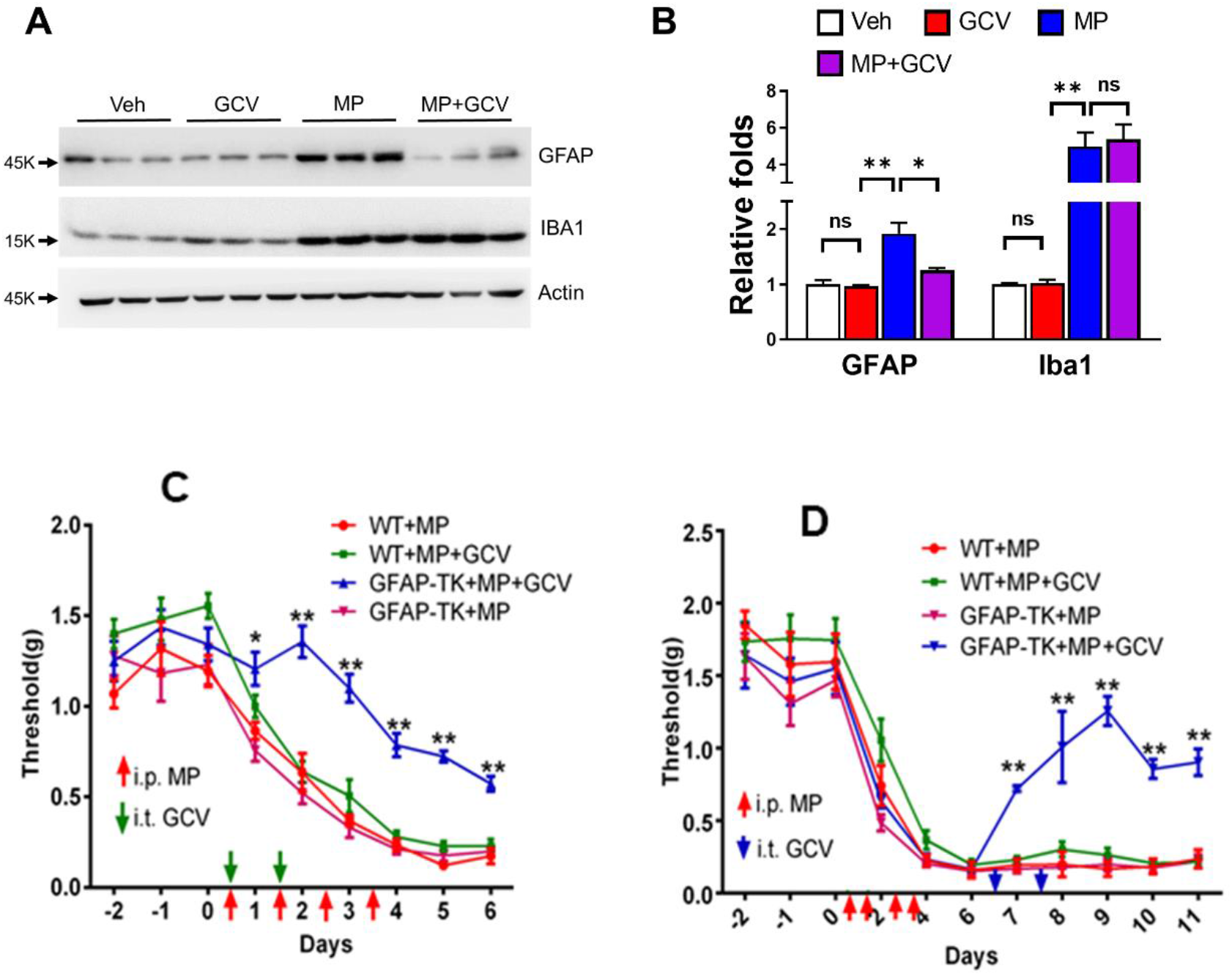
Reactive astrocytes are critical for morphine-induced hyperalgesia. (**A**) Immunoblotting analysis of GFAP and Iba1 in the spinal cord of the GFAP-TK transgenic mice treated with GCV, morphine or GCV+morphine using the drug administration paradigm shown in C. Mice were euthanized for spinal cord collection at day 7. (**B**) Quantitative summary of band intensity of immunoblots. GCV abolished morphine-induced up-regulation of GFAP but not Iba1, indicating GCV ablated specifically reactive astrocytes but not reactive microglia in GFAP-TK mice. GFAP and Iba1 protein levels were determined by immunoblotting analysis. *p<0.05; **p<0.01; ns, p>0.05. (**C**) Effect of astrogliosis ablation on OIH. GFAP-TK mice expressed mechanical OIH similarly as WT mice. Injections of GCV (green arrows; 5 mg/kg; i.t.) during the early phase of morphine (MP) administration (red arrows; 20 mg/kg; i.p.) inhibited the morphine-induced increase of mechanical sensitivity, measured by von Frey tests, specifically in GFAP-TK transgenic but not in WT mice (n=6/group). (**D**) Effect of astrogliosis ablation on OIH maintenance. Injections of GCV (blue arrows; i.t.) after OIH was established by morphine administration (red arrows; i.p.) reversed the morphine-induced increase of mechanical sensitivity in GFAP-TK transgenic but not in WT mice. (n=6/group).

In these animal models, we found microglia and astrocyte activation in the spinal dorsal horn (SDH) (Fig. 1A-1B), consistent with previous observations^35, 36^. A critical role of microglia in OIH was proposed in an early study^11^, but inconsistent findings were reported later^4, 12^. In this study, we aimed to examine the potential role of reactive astrocytes in OIH development. To this end, we employed a genetic approach to selectively ablate reactive astrocytes, using GFAP-thymidine kinase (TK) transgenic mice^37^. The transgenic mice were administered with ganciclovir (GCV), which would be metabolized to nucleotide analogues by TK in astrocytes. The GCV-derived nucleotide analogues are toxic to proliferating cells, and thus would specifically induce cell death of reactive astrocytes.

Von Frey tests showed that the GFAP-TK transgenic mice expressed mechanical OIH similarly as WT mice (Fig. 1C). To determine the contribution of astrogliosis to the expression of OIH, we intrathecally (i.t.) injected GCV (5 mg/kg) during the first 2 days of morphine administration. We observed that GCV administration blocked the morphine-induced increase of GFAP but not Iba1 (Fig. 1A–1B), indicating selective ablation of reactive astrocytes without affecting microglia. Importantly, the GCV treatment impaired OIH expression in GFAP-TK but not in WT mice (Fig. 1C). Significant GCV-induced impairment of OIH in the transgenic mice started to develop one day after the first GCV injection, suggesting a critical role of reactive astrocytes in early expression of OIH.

We then sought to determine if astrogliosis contributes to OIH maintenance. To this end, we tested the effect of ablation of reactive astrocytes on established OIH. In this experiment, GCV was administered six days after the first injection of morphine, when OIH was fully expressed (Fig. 1D). We observed that the GCV administration significantly reversed the established OIH in GFAP-TK but not in WT mice (Fig. 1D), indicating an important role of reactive astrocytes in OIH maintenance. These results collectively showed that reactive astrocytes were essential for both expression and maintenance of OIH.

To test potential sex dimorphism of the astrocytic role in OIH, we determined the effect of selective astrocytic toxin L-AA (i.t., 100nmol/5μ on days 0, 2, 4, 6) OIH expression induced in female mice. We observed that L-AA treatment significantly inhibited OIH (Supplemental Fig 2), indicating astrocytes also play crucial for OIH development in female mice.

### Reactive astrocytes mediate morphine-induced neural circuit polarization in the SDH

To gain insight into the neural circuitry mechanism by which astrogliosis contributes to OIH, we tested the effect of astrogliosis ablation on morphine-induced malplasticity in pain neural circuits. When we performed whole-cell recording of SDH neurons in morphine-treated GFAP-TK mice, we found that morphine increased both the frequency of spontaneous excitatory postsynaptic currents (sEPSCs) and the amplitude of evoked EPSCs (eEPSCs) of excitatory neurons (Fig. 2A–2C), identified by their characteristic non-tonic firing patterns ^38, 39^. By contrast, morphine decreased the sEPSC frequency and eEPSC amplitude of inhibitory neurons (Fig. 2D–2F), identified by their characteristic tonic firing pattern ^38, 39^. These findings indicate that morphine administration polarizes neural circuits in the SDH, by increasing excitatory inputs to excitatory neurons and meanwhile decreasing excitatory inputs to inhibitory neurons. The observed neural circuit polarization (NCP) might contribute to OIH expression, by facilitating activation of the SDH pain pathway.

**Figure 2.**
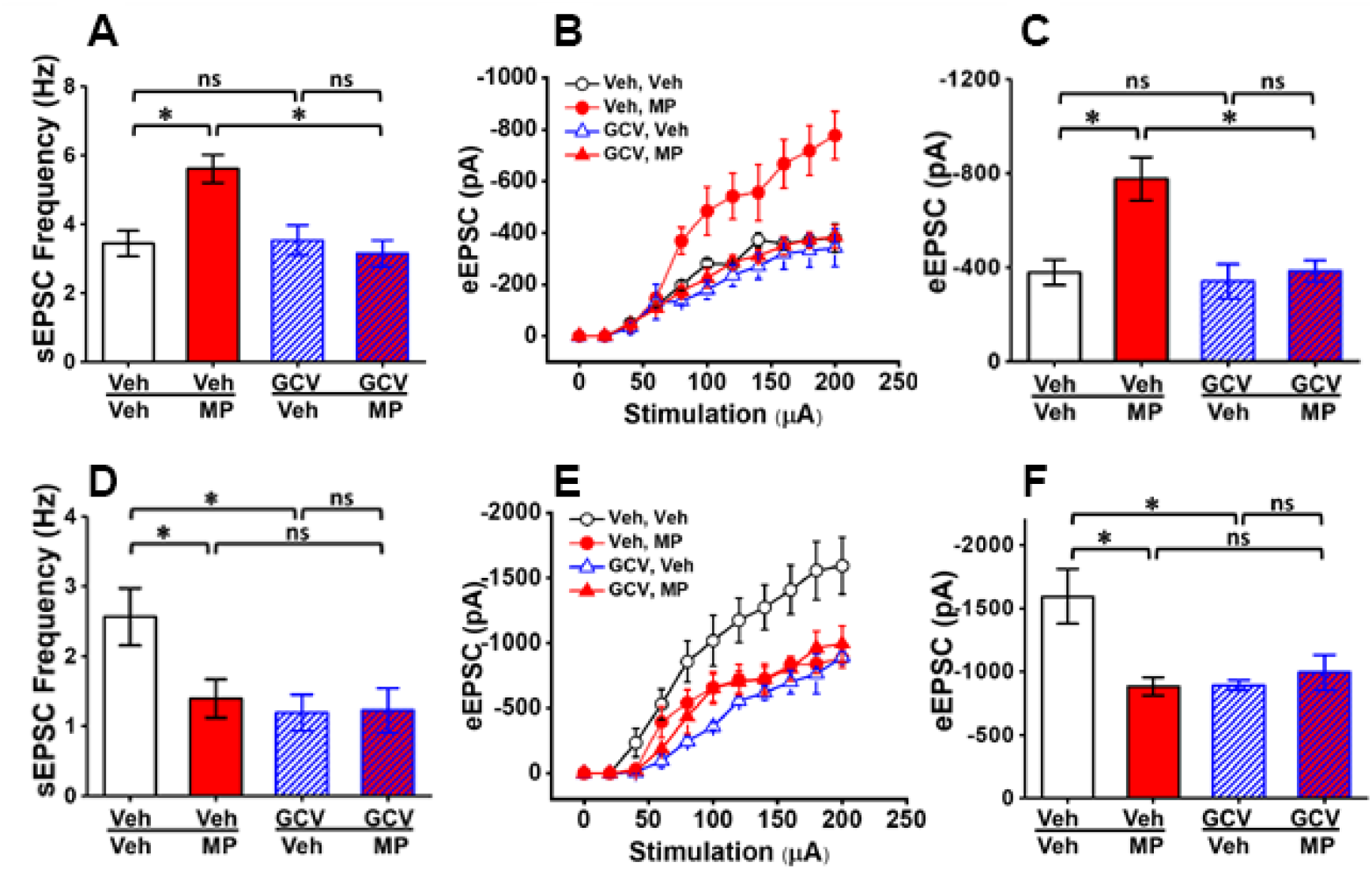
Reactive astrocytes are critical for morphine-induced NCP in the SDH. (**A**) Effect of astrocytic ablation on sEPSCs of non-tonic firing neurons in the SDH of GFAP-TK transgenic mice. Whole-cell patch recording was performed from spinal slices prepared from GFAP-TK transgenic mice (day 7) treated with morphine and/or GCV according to the paradigm shown in Fig. 1C. Morphine increased sEPSC frequency, whereas GCV abolished this increase. GCV did not affect basal sEPSC frequency. Veh/Veh: 21/3 (cells/mice); veh/MP: 60/5; GCV/Veh: 29/4; GCV/MP: 50/5. (**B**) Effect of astrogliosis ablation on eEPSCs of non-tonic firing neurons in the SDH of GFAP-TK transgenic mice. eEPSC amplitudes of patched SDH neurons were recorded from spinal slices prepared as in A. Morphine increased eEPSC amplitude, whereas GCV abolished this increase. GCV did not affect basal eEPSC amplitude. Veh/Veh: 24/3 (cells/mice); veh/MP: 48/4; GCV/Veh: 35/3; GCV/MP: 48/4 (**C**) Statistical analysis of eEPSC amplitude evoked by 200 µA stimulation shown in B. (**D**) Effect of astrogliosis ablation on sEPSCs of tonic firing neurons in the SDH of GFAP-TK transgenic mice. Morphine decreased sEPSC frequency. GCV alone also decreased sEPSC frequency. However, morphine did not further decrease sEPSC frequency after GCV treatment. Veh/Veh: 13/3 (cells/mice); veh/MP: 12/4; GCV/Veh: 20/5; GCV/MP: 26/5. (**E**) Effect of astrogliosis ablation on eEPSCs of tonic firing neurons in the SDH of GFAP-TK transgenic mice. Morphine decreased eEPSC amplitude. GCV alone also decreased eEPSC amplitude. However, morphine did not further decrease eEPSC amplitude after GCV treatment. Veh/Veh: 14/3 (cells/mice); veh/MP: 12/4; GCV/Veh: 20/3; GCV/MP: 26/4. (**F**) Statistical analysis of eEPSC amplitude evoked by 200 µA stimulation shown in E. *p<0.05; **p<0.01; ns, p>0.05.

Astrocytes normally play important role in maintaining homeostasis of neural circuits ^17^, but it is unclear how astrogliosis modulates neural circuits during pain pathogenesis. Hence, we set to test the contribution of astrogliosis to morphine-induced NCP. To this end, we determined the effect of astrogliosis ablation on EPSCs of excitatory and inhibitory neurons in the SDH of GFAP-TK transgenic mice. We found that GCV treatment abolished the morphine-induced increase in sEPSC frequency and eEPSC amplitude of excitatory neurons (Fig. 2A–2C).

On inhibitory neurons, we observed that GCV treatment by itself decreased sEPSC frequency and eEPSC amplitude of inhibitory neurons in GFAP-TK mice without morphine administration (Fig. 2D-2F), indicating excitatory inputs on inhibitory neurons under physiological conditions are more sensitive to astrocyte ablation than excitatory inputs on excitatory neurons. Importantly, after GCV treatment morphine failed to induce further decease of sEPSC frequency and eEPSC amplitude on inhibitory neurons (Fig. 2D-2F). Together, these results indicated that astrogliosis was critical for morphine to induce NCP, including the increase of sEPSC frequency and eEPSC amplitude on excitatory neurons and the decrease of sEPSC frequency and eEPSC amplitude on inhibitory neurons.

### Neuronal Wnt5a is critical for morphine to induce astrogliosis, NCP and OIH

Having shown the essential role of astrogliosis in OIH and NCP, we next sought to elucidate the mechanism by which morphine induces astrocyte activation. To this end, we focused on the potential role of Wnt5a signaling, which is implicated in pain pathogenesis^40–42^. Wnt5a is secreted protein mainly expressed by neurons^43^, and is upregulated by in the SDH by pain signals^44^. Wnt5a secretion is controlled by synaptic activity^45, 46^. The involvement of Wnt5a in regulating astrogliosis is suggested by the observations that Wnt5a antagonist inhibits astrocyte activation in pain models^47, 48^ and JNK, a critical downstream target in the Wnt5a signaling pathway, is critical for astrogliosis in a neuropathic pain model^49^. Based on these prior findings, we hypothesized that Wnt5a secreted from neurons hyperactivated following morphine treatment activated astrocytes. To test this hypothesis, we generated neuronal Wnt5a conditional knock-out (CKO) mice (Wnt5a-CKO-S) by crossing floxed Wnt5a mice^22^ with synapsin 1-Cre mice^23^. We observed that morphine administration significantly upregulated spinal Wnt5a protein in the WT mice in a time-dependent pattern, which peaked at day 3 (Supplemental Fig 3). However, Wnt5a-CKO-S mice did not show morphine-induced Wnt5a protein upregulation (Fig. 3A), indicating morphine stimulated Wnt5a expression predominantly in neurons. The morphine-induced Wnt5a increase was inhibited by NMDAR antagonist APV (i.t., 5μg/5μl, 30 min prior to morphine injection) (Supplemental Fig 4), indicating NMDAR activation mediated morphine-induced Wnt5a protein upregulation, as suggested by previous studies^45^. Importantly, we found that, unlike WT mice, morphine completely failed to induce GFAP upregulation in the spinal cord of the Wnt5a-CKO-S mice (Fig. 3B), indicating deletion of Wnt5a in neurons abolished morphine-induced astrogliosis. To confirm these findings, we also generated another Wnt5a CKO mutant to delete Wnt5a in neural stem cells using nestin-Cre^24^ (Wnt5a CKO-N). We found that this Wnt5a mutation also abolished morphine-induced astrogliosis (Supplemental Fig 5). These findings collectively showed that neuronal Wnt5a was essential for morphine to induce astrogliosis.

**Figure 3.**
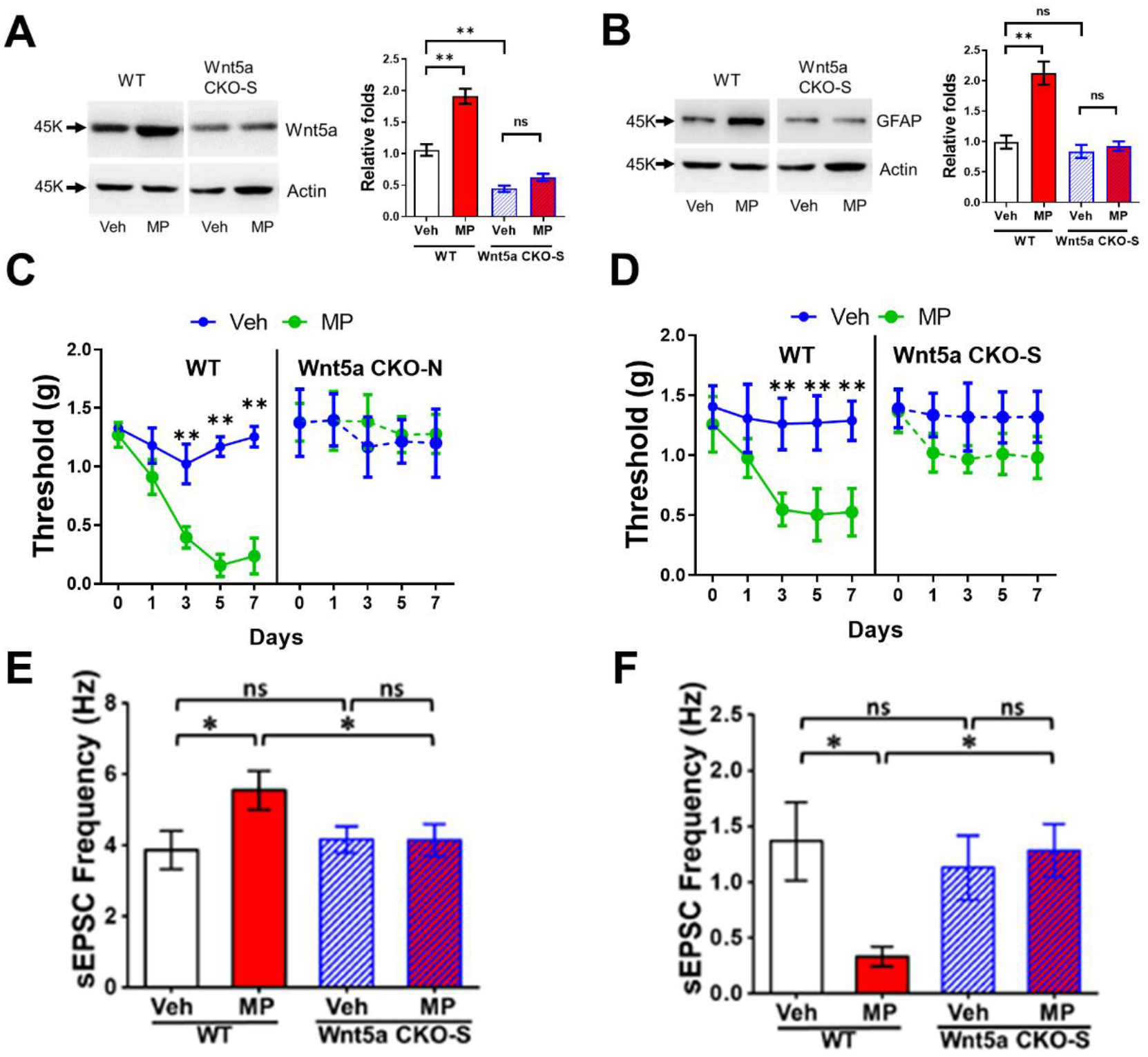
Neuronal Wnt5a is essential for morphine-induced astrogliosis, hyperalgesia, and NCP in the SDH. Two Wnt5a CKO mouse lines were generated, one by crossing floxed Wnt5a with nestin-Cre mice (Wnt5a CKO-N) and the other by crossing floxed Wnt5a with synapsin 1-Cre mice (Wnt5a CKO-S). (**A**) Wnt5a CKO-S (and Wnt5a CKO-N, not shown) blocked morphine (MP)-induced upregulation of Wnt5a in the spinal cord. (**B**) Wnt5a CKO-S (and Wnt5a CKO-N, Sup. Fig. 2) blocked morphine-induced spinal astrogliosis, as shown by GFAP upregulation. Tissues used in **A** and **B** were collected from mice (n=4/group) on day 7 after morphine administration according to the paradigm shown in Fig. 1A. (**C**) Wnt5a CKO-S abolished OIH expression (n=6/group). (**D**) Wnt5a CKO-N abolished OIH expression (n=6/group). (**E**) Wnt5a CKO-S blocked the morphine-induced increase in sEPSC frequency of non-tonic firing neurons in the SDH. Veh/WT, 19/3 (cells/mice); MP/WT: 47/5; Veh/Wnt5a CKO-S, 57/5; MP/Want5a CKO-S, 33/4. (**F**) Wnt5a CKO-S blocked the morphine-induced decrease in sEPSC frequency of tonic firing neurons in the SDH. Veh/WT, 12/3 (cells/mice); MP/WT: 18/3; Veh/Wnt5a CKO-S, 28/5; MP/Want5a CKO-S, 13/4.

As reactive astrocytes were crucial for OIH induced by morphine (Fig. 1C, 1D) and neuronal Wn5a was required for the astrocyte activation (Fig. 3B), we hypothesized that the OIH expression depended on neuronal Wnt5a. To test this hypothesis, we determined the effect of neuronal Wnt5a CKO on OIH. The result showed that OIH expression was abolished in Wnt5a CKO-S mice (Fig. 3C). We observed similar inhibitory effect on OIH in Wnt5a CKO-N mice (Fig. 3D). These data together demonstrated the requirement of neuronal Wnt5a for OIH development.

Because of the observed role of astrogliosis in NCP (Fig. 2) and neuronal Wnt5a in the astrogliosis (Fig. 3B), we further hypothesized that neuronal Wnt5a was important for NCP expression. To test this, we determined the expression of NCP in the SDH of Wnt5a CKO-S mice. We found that the morphine-induced increase of sEPSC frequency of excitatory neurons (Fig. 3E) and the decrease of sEPSC frequency of inhibitory neurons (Fig. 3F) were abolished in Wnt5a CKO-S mice. These results showed that neuronal Wnt5a mediated the expression of NCP in the SDH.

### Astrocytic ROR2 receptor is crucial for astrogliosis, NCP and OIH induced by morphine

The results described above demonstrate a key role of Wnt5a from neurons in mediating morphine-induced astrocyte activation. Next, we sought to further elucidate the underlying mechanism by which Wnt5a activates astrocytes. We hypothesized that Wnt5a co-receptor ROR2 on astrocytes ^43^ was critical for the astrogliosis. To test this hypothesis, we generated astrocytic CKO of ROR2 by crossing floxed ROR2^25^ and GFAP-Cre^26^ mouse lines. Similar to the effect of neuronal Wnt5a CKO (Fig. 3), we observed that the astrocytic ROR2 CKO also blocked morphine-induced GFAP upregulation, indicating inhibition of astrogliosis (Fig. 4A).

**Figure 4.**
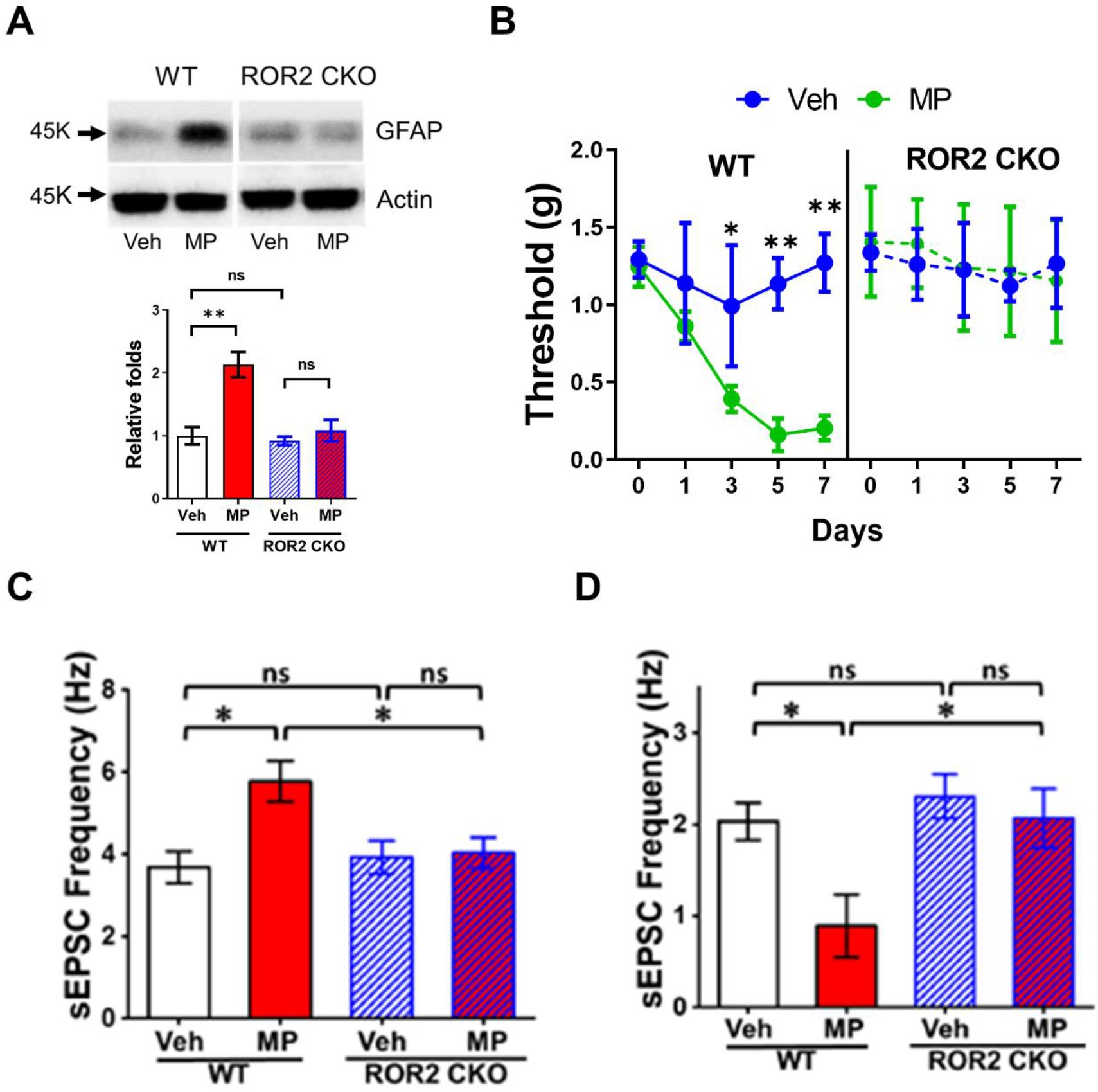
Astrocytic ROR2 is required for morphine-induced astrogliosis, hyperalgesia, and NCP in the SDH. (**A**) Astrocytic ROR2 CKO blocked morphine-induced astrogliosis. Spinal cords were collected from mice (n=4/group) on day 7 after morphine administration according to the paradigm shown in Fig. 1A. (**B**) Astrocytic ROR2 CKO blocked OIH expression (n=6/group). (**C**) Astrocytic ROR2 CKO blocked the morphine-induced increase in sEPSC frequency of non-tonic firing neurons in the SDH. Veh/WT, 32/3 (cells/mice); MP/WT: 53/5; Veh/ROR2 CKO, 45/3; MP/ROR2 CKO, 47/4. (**D**) Astrocytic ROR2 CKO blocked the morphine-induced decrease in sEPSC frequency of tonic firing neurons in the SDH. Veh/WT, 19/3 (cells/mice); MP/WT: 18/3; Veh/ROR2 CKO, 22/3; MP/ROR2 CKO, 12/3.

As astrogliosis and neuronal Wnt5a are critical for OIH development (Fig. 1; Fig. 3), the finding of the astrogliosis blockage by the ROR2 CKO led us to predict that the mutant mice would be impaired in OIH development. In support of this hypothesis, we indeed found that the expression of OIH was abolished in the ROR2 CKO mice (Fig. 4B), as in the Wnt5a CKO-S and Wnt5a CKO-N mice (Fig. 3).

Because of the observed contribution of astrogliosis and neuronal Wnt5a to NCP (Fig. 2; Fig. 3), we further hypothesized that the ROR2 CKO would impair the expression of NCP. We tested this hypothesis by determining effect of the ROR2 CKO on sEPSCs of neurons in the SDH. The results showed that both the morphine-induced increase in sEPSC frequency of excitatory neurons and decrease in sEPSC frequency of inhibitory neurons in the SDH were blocked in ROR2 CKO mice (Fig. 4C-4D). The findings suggested that, similar to neuronal Wnt5a, astrocytic ROR2 also played an important role in morphine-induced astrogliosis, OIH, and NCP.

### Wnt5a signaling-regulated reactive astrocytes control OIH and NCP via IL-1β

The above results from Wnt5a CKO and ROR2 CKO mice collectively suggest that Wnt5a secreted from neurons stimulates astrocytic ROR2 receptor to promote morphine-induced astrogliosis, and that this neuron-to-astrocyte Wnt5a-ROR2 signaling pathway contributes to OIH expression via astrogliosis. Next, we sought to further understand the mechanism by which Wnt5a signaling-mediated astrogliosis regulates OIH pathogenesis. Previous studies reveal that reactive astrocytes contribute to pain pathogenesis by releasing proinflammatory mediators^16^. Indeed, we observed morphine-induced upregulation of spinal activated (cleaved) IL-1β but not pro-IL-1β (Fig. 5A), a key proinflammatory cytokine that is implicated in OIH^50^ and regulated by Wnt5a signaling^43^. Importantly, CKO of Wnt5a in either Wnt5a CKO-S or CKO-N mice abolished the IL-1β upregulation, without affecting pro-IL-1β levels (Fig. 5A). CKO of ROR2 in astrocytes also blocked upregulation of active IL-1β (Fig. 5A). These results suggested that the neuron-to-astrocyte Wnt5a-ROR2 signaling controlled morphine-induced IL-1β upregulation.

**Figure 5.**
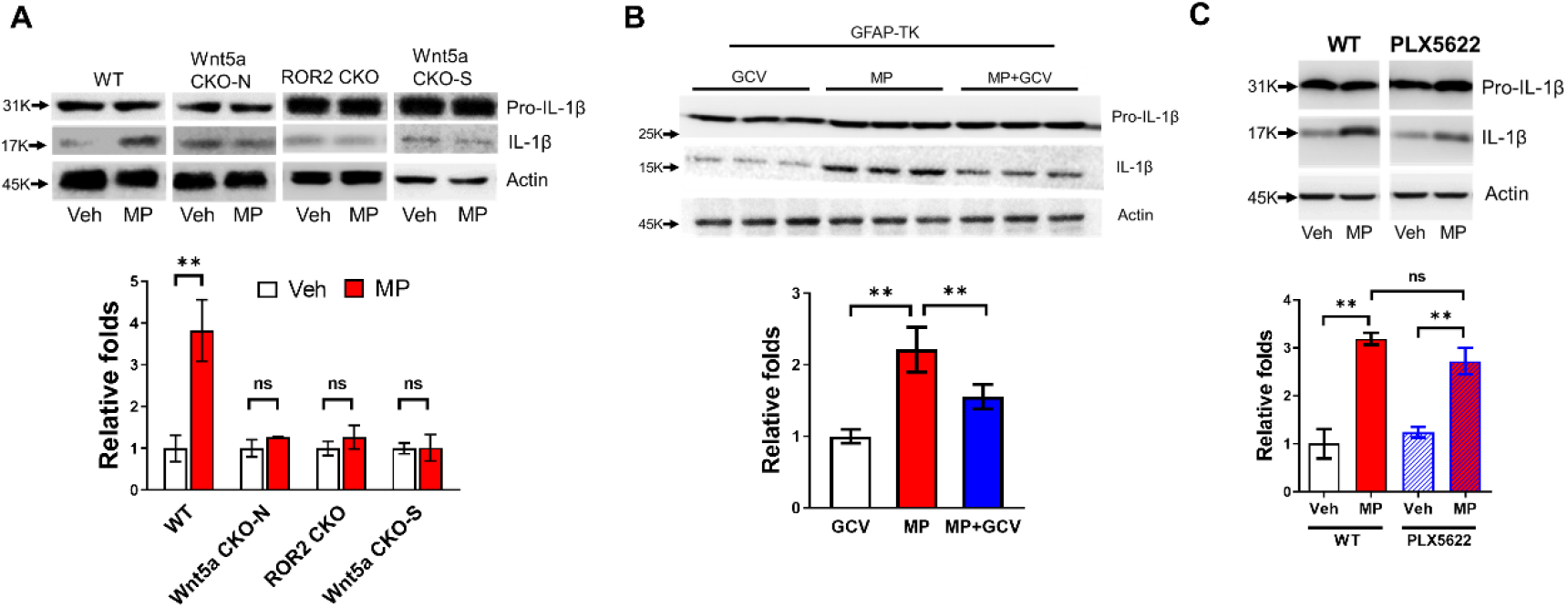
Morphine induces IL-1β activation (cleavage) via Wnt5a-ROR2 signaling-dependent astrogliosis. (**A**) Neuronal Wnt5a CKO and astrocytic ROR2 CKO blocked morphine-induced IL-1β upregulation without affecting pro-IL-1β. Spinal cords were collected for immunoblotting from mice (n=4/group) on day 7 after morphine administration according to the paradigm shown in Fig. 1A. (**B**) Astrogliosis ablation in GFAP-TK mice inhibited morphine-induced spinal IL-1β upregulation as revealed by immunoblotting (n=4/group). (**C**) Microglial ablation by PLX5622 did not affect morphine-induced spinal IL-1β upregulation as revealed by immunoblotting (n=4/group).

Because the neuron-to-astrocyte Wnt5a-ROR2 signaling controls both astrocyte activation (Fig. 3; Fig.4) and IL-1β upregulation induced by morphine (Fig. 5A), we hypothesized that the reactive astrocytes produced IL-1β. To test this hypothesis, we determined the effect of ablating reactive astrocytes on the IL-1β upregulation. We found that astrogliosis ablation by GCV in GFAP-TK transgenic mice significantly impaired morphine-induced IL-1β upregulation in the spinal cord (Fig. 5B). On the other hand, ablation of microglia by the CSF1R inhibitor PLX5622^51–53^ did not significantly affect morphine-induced spinal IL-1β upregulation (Fig. 5C). These results suggested that reactive astrocytes, rather than microglia, were the major cell source of morphine-induced IL-1β. Previous studies suggest astrocytic IL-1β in other pain models^13^.

Because reactive astrocytes contributed to morphine-induced OIH (Fig. 1) and produced IL1β (Fig. 5B), we hypothesized that astrogliosis promoted OIH via IL-1β. Hence, we tested the pathogenic role of IL-1β in OIH expression. In these experiments, we used the endogenous IL-1 receptor antagonist IL-1Ra to block Il-1β signaling ^50^. We observed that IL-1Ra administration (20 μg/kg/day for the first 3 days, i.t.) abolished the expression of OIH, without affecting baseline mechanical sensitivity (Fig. 6A). These results suggested that IL-1β produced by reactive astrocytes was critical for OIH pathogenesis.

**Figure 6.**
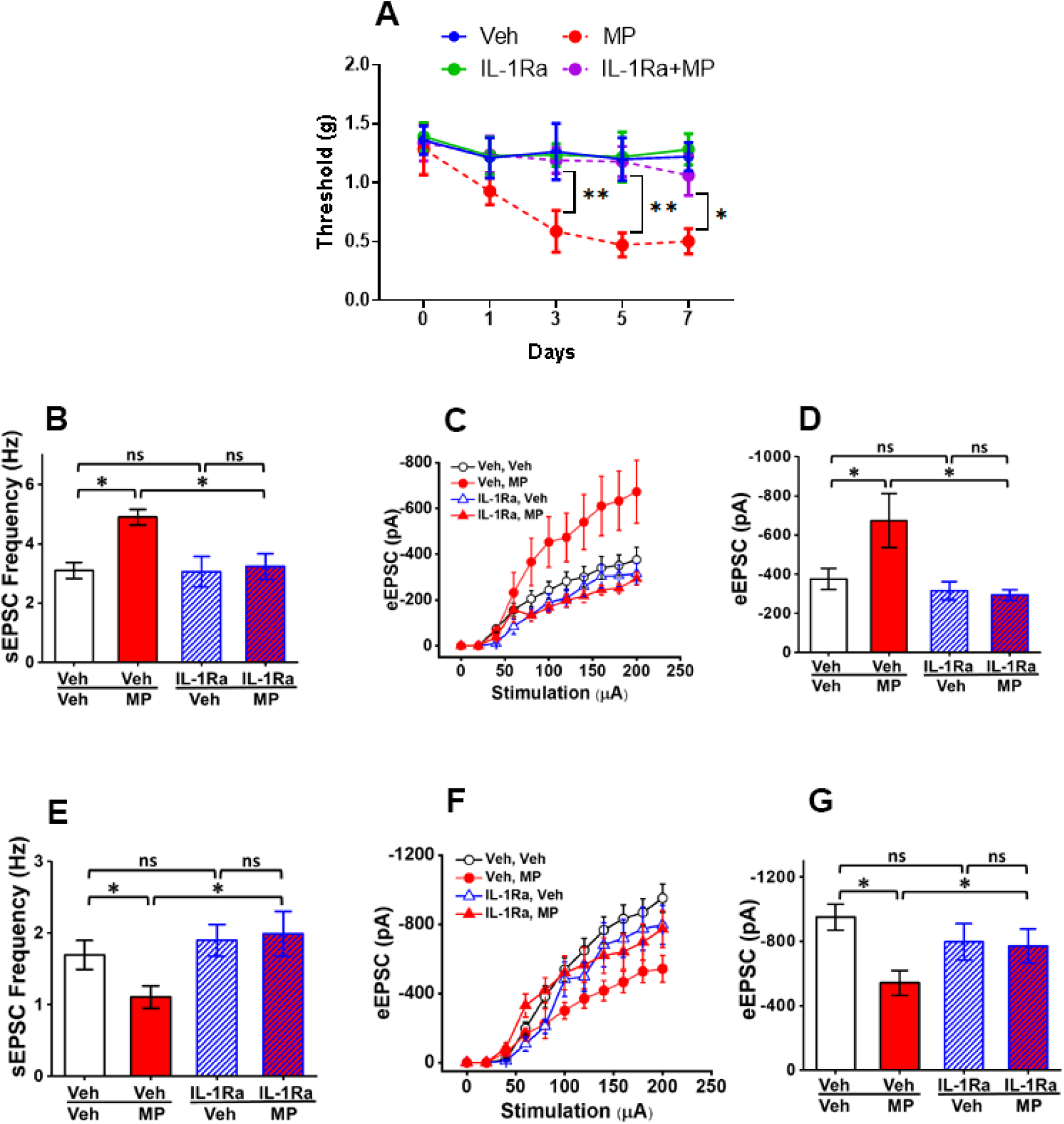
IL-1Ra blocks OIH and NCP. (**A**) The IL-1β receptor antagonist IL-1Ra blocked OIH expression. Morphine was administered according to the paradigm in Fig. 1C, and IL-1Ra was administered daily on the first 3 days (20 μg/kg, i.t.). (**B**) IL-1Ra impaired the morphine-induced increase in sEPSC frequency of non-tonic firing SDH neurons (Veh/Veh: 24/3 (cells/mice); veh/MP: 33/5; IL-1Ra/Veh: 22/5; IL-1Ra/MP: 44/4) in spinal slices prepared on day 7 from mice that received drug administration as infIG. 2A. (**C, D**) IL-1Ra abolished the morphine-induced increase in eEPSC amplitude of non-tonic firing neurons in the SDH; D shows statistical analysis of eEPSC amplitudes evoked by 200 µA stimulation shown in C (Veh/Veh: 24/3 (cells/mice); veh/MP: 28/5; IL-1Ra/Veh: 19/3; IL-1Ra/MP: 40/5). (**E**) IL-1Ra blocked the morphine-induced decrease in sEPSC frequency of SDH inhibitory neurons from GAD67-GFP transgenic mice (Veh/Veh: 67/3 (cells/mice); veh/MP: 51/5; IL-1Ra/Veh: 35/5; IL-1Ra/MP: 44/4). (**F, G**) IL-1Ra abolished the morphine-induced decrease in eEPSC amplitude of SDH inhibitory neurons; G shows statistical analysis of eEPSC amplitudes evoked by 200 µA stimulation shown in I (Veh/Veh: 42/4 (cells/mice); veh/MP: 43/5; IL-1Ra/Veh: 31/3; IL-1Ra/MP: 39/4). *p<0.05; **p<0.01; ns, p>0.05.

IL-1Ra administration also blocked the morphine-induced increase in sEPSC frequency (Fig. 6B) and eEPSC amplitude (Fig. 6C-6D) in excitatory neurons in the SDH. Furthermore, using GAD67-GFP transgenic mice^54^, we found that IL-1Ra blocked the morphine-induced decrease in sEPSC frequency (Fig. 6E) and sEPSC amplitude (Fig. 6F-6G) of GABAergic inhibitory neurons in the SDH. Together, these results indicated that the morphine-induced expression of NCP was controlled by IL-1β produced by Wnt5a-ROR2 signaling-activated astrocytes (Fig. 5A-5B).

### Wn5a-ROR2 signaling activates IL-1β via Inflammasomes

Having identified a critical role of IL-1β in OIH, we wanted to further understand the mechanism by which morphine regulates astrocytic IL-1β. We hypothesized inflammasome, a key protein complex that controls IL-1β processing^55^ and is critical for OIH^56^ and for morphine to prolong neuropathic pain^57^, played a key role in this process. We observed that morphine administration upregulated the active forms of both IL-1β and caspase 1 (Cas-1), the inflammasome proteinase that processes pro-IL-1β protein, but, interestingly, did not affect the expression of pro-IL-1β and pro-caspase 1 (Fig. 7A) and their mRNA (Supplemental Fig 6). In addition, pharmacological blockade of the inflammasome with AC-YVAD-CMK, a selective inhibitor of Cas-1, impaired the upregulation of cleaved Cas-1 and IL-1β without compromising pro-IL-1β and pro-caspase 1 protein expression (Fig. 7A). These results indicated that morphine stimulated spinal IL-β activation via inflammasome activation. Furthermore, inhibition of the inflammasome by AC-YVDA-CMK or Cas-1 siRNA also blocked the expression of OIH (Fig. 7B-7C). These data collectively demonstrated a critical role of inflammasome in the regulation of IL-1β during OIH expression.

**Figure 7.**
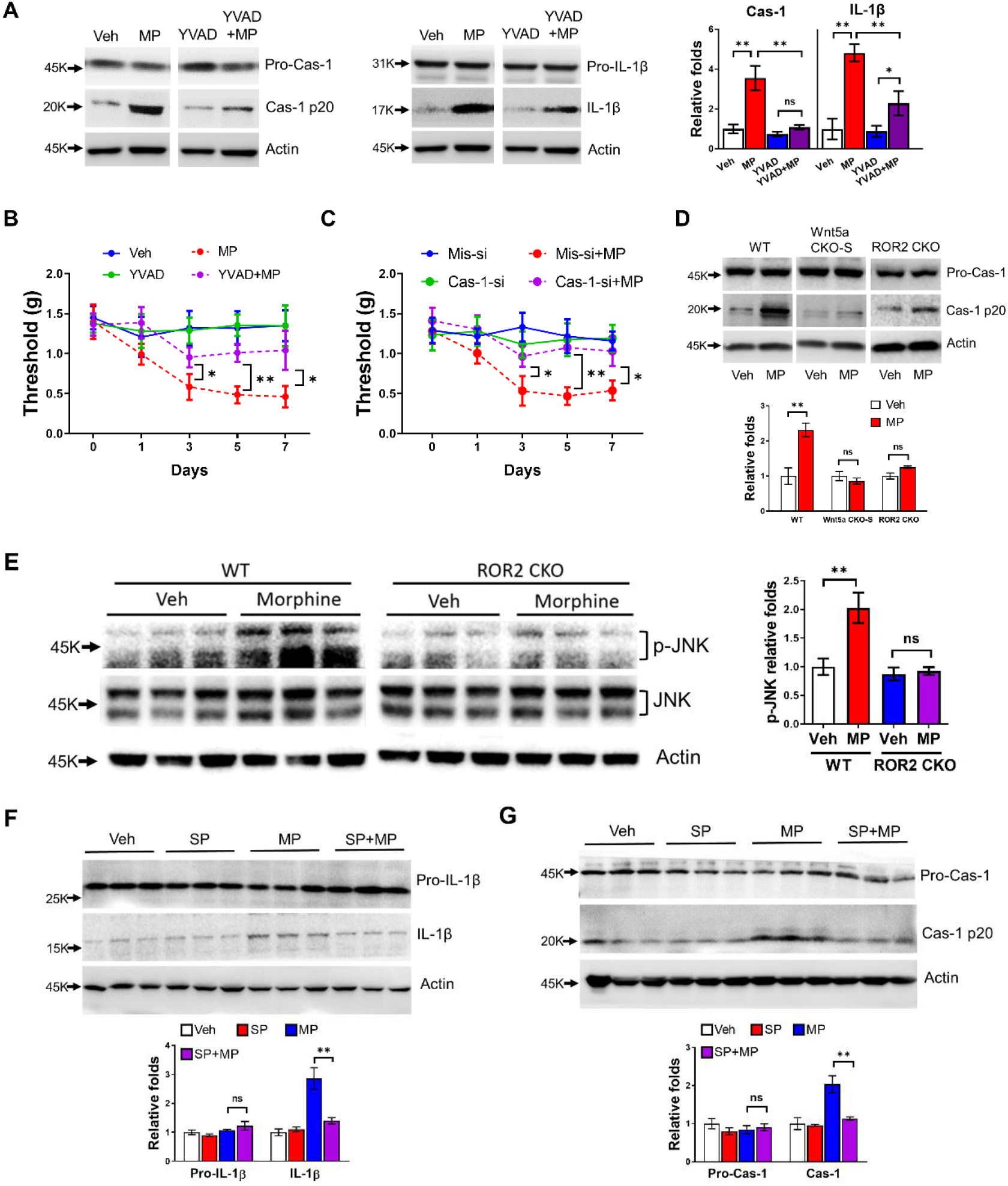
The Wnt5a-ROR2 signaling pathway controls inflammasome activation during OIH development. (**A**) Morphine activated the inflammasome in the spinal cord as measured by the upregulation of Cas-1 and IL-1β. Morphine-induced inflammasome activation was impaired by AC-YVAD-CMK (YVAD), a selective inhibitor of Cas-1. (**B**) YVAD impaired the expression of mechanical OIH. Morphine was administered according the paradigm shown in Fig. 1A, and YVAD was administered daily on the first 4 days (4 nmol/kg, i.t.). (**C**) Cas-1 siRNA (Cas-si) but not mismatch (silencer negative control) siRNA (Mis-si) blocked mechanical OIH expression. siRNA was administered daily on the first 4 days (3 nmol/kg, i.t.). (**D**) Morphine-induced inflammasome activation, measured by Cas-1 immunoblotting, was blocked by neuronal Wnt5a-CKO-S or astrocytic ROR2 CKO. (**E**) Immunoblotting analysis of spinal phosphorylated and total JNK in wild type or ROR2 CKO mice treated with morphine according to the paradigm shown in Fig. 4A. Morphine-induced phosphorylation of JNK were blocked by astrocytic ROR2 CKO. (**F**) Immunoblotting analysis of pro-IL-1β and IL-1β in the spinal cord of wild type mice treated with saline, SP600125, morphine or SP+morphine. SP was intrathecally administered (10μg/5μl) at 0,2,4,6 day and 30 minutes prior to morphine injection. Mice were euthanized for spinal cord collection at day 7 (n=4/group). Morphine-induced IL-1β activation was blocked by p-JNK inhibitor SP600125. (**G**) Immunoblotting analysis of Pro-Cas-1 and Cas-1 in the spinal cord of wild type mice treated with saline, SP600125, morphine or SP+morphine. Morphine-induced Cas-1 activation was blocked by p-JNK inhibitor SP600125. *p<0.05; **p<0.01; ns, p>0.05.

Because morphine induced IL-1β via Wnt5-ROR2 signaling-regulated astrogliosis (Fig. 3; Fig. 4; Fig. 5B), we hypothesized the neuron-to-astrocyte Wnt5a-ROR2 signaling regulated morphine-induced inflammasome activation. To test this idea, we determined the effect of neuronal Wnt5a CKO and astrocytic ROR2 CKO on inflammasome activation, by measuring Cas-1 levels. We found that either the Wnt5a CKO-S or ROR2 CKO abolished the morphine-induced increase of active Cas-1, without significant change of pro-Cas-1 levels (Fig. 7D). These findings suggested that Wnt5a-ROR2 signaling mediated morphine-induced inflammasome activation to control IL-1β maturation in astrocytes.

The observations that morphine induced upregulation of cleaved Cas-1 and IL-1β (Fig. 7A) without changes of pro-Cas-1 (Fig. 7A, 7D) and pro-IL-1β levels (Fig. 5A-C; Fig. 7A) indicate that in the OIH model morphine induces mainly inflammasome activation, rather than priming. To gain insights into the mechanism by which Wnt5a-ROR2 signaling controls morphine-induced inflammation activation, we sought to test the role of JNK, because JNK is a key downstream target of ROR2^58^ and plays critical role in priming/activation of inflammasome^59^. We found that active (phosphorylated) JNK was upregulated by morphine in wild-type mice, and astrocytic ROR2 CKO inhibited morphine-induced JNK activation (Fig. 7E). These data suggest that morphine elicits JNK activation via astrocytic ROR2 in the spinal cord. To test the role of JNK in inflammasome activation induced by morphine, we determined the effects of intrathecal administration of JNK selective inhibitor SP600125 (i.t., 10μg/5μl, 30 min before morphine injection) on the levels of IL-1b and Casp1, as well as pro-IL-1b and pro-casp-1. We found that the JNK inhibitor blocked morphine-induced increase of cleaved IL-1β (Fig. 7F) and Cas-1 (Fig. 7G) without affecting their precursors (Fig. 7F-7G), indicating JNK is critical for morphine-induced inflammasome activation. These data together suggest activation of ROR2-JNK signaling induced by morphine leads to inflammasome activation.

## DISCUSSION

We show that astrogliosis is crucial for morphine-induced OIH and expression of NCP in the SDH. The morphine-induced astrogliosis, NCP, and OIH requires Wnt5a in neurons and the ROR2 receptor in astrocytes. Furthermore, the effects of morphine on neural circuits and pain behavior are mediated by IL-β controlled by Wnt5a-ROR2 pathway-regulated inflammasome activation. These findings suggest bidirectional interactions between neurons and astrocytes during the pathogenesis of OIH. We propose that morphine induces Wnt5a release from neurons to stimulate astrocytic ROR2 receptor to activate astrocytes and inflammasome to produce IL-1β, and active IL-1β released from reactive astrocytes in return stimulates pain processing neurons in the SDH to promote NCP and OIH (Fig. 8).

**Figure 8.**
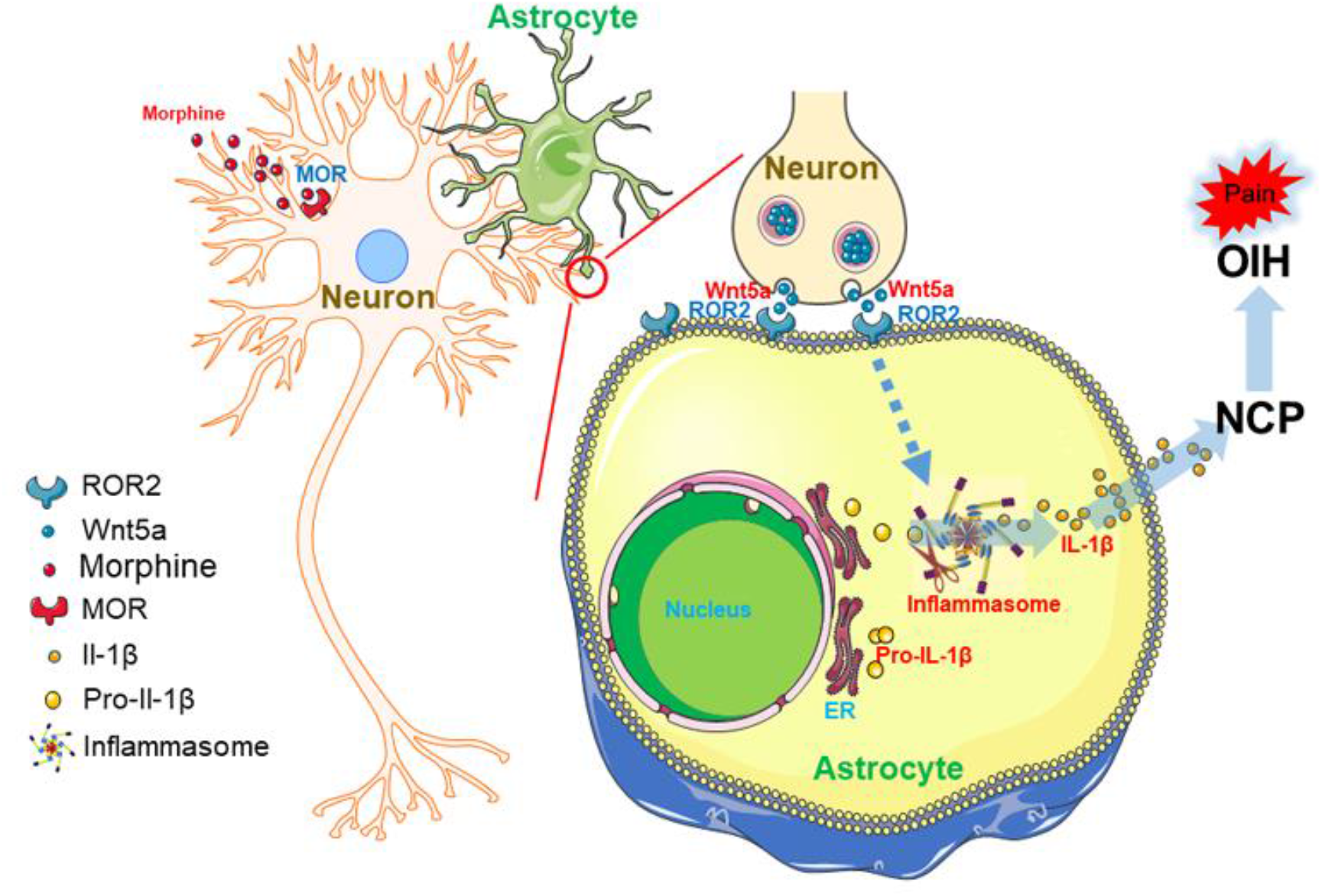
Model of the neuron-to-astrocyte Wnt5a-ROR2 signaling pathway in OIH development. Wnt5a-ROR2 signaling controls morphine-induced astrogliosis and astrocytic IL-1β activation. IL-1β is activated by the inflammasome, and feedback to neurons induces OIH.

To evaluate the potential relevance of our results from mouse models to human conditions, we performed snRNA-seq analysis of human lumbar spinal cords and compared their neuron and astrocyte subtypes with that of mouse spinal cords. We found that, the human neurons^31^ and astrocytes^32^ and the mouse counterparts showed similar clusters based on the transcriptomic characteristics, suggesting the similar heterogeneity in neurons and astrocytes of human and mouse spinal cords. In addition, we compared the expression profiles of Wnt5a in neurons and ROR2 in astrocytes, and observed that Wnt5a was expressed in homologous neuronal clusters (Suppl. Fig.7) and ROR2 is expressed in the homologous astrocyte clusters of human and mouse spinal cords (Suppl. Fig. 8). These findings indicate similar neuronal and astrocytic heterogeneity and similar cellular expression profiles of Wnt5a and ROR2 in neurons and astrocytes respectively in the human and mouse spinal cords.

### Neuronal circuitry mechanism in the development of OIH

Malplasticity of multiple components in the pain transmission pathway is implicated in OIH. Sensitization of peripheral nociceptors and second-order neurons in the spinal cord, long-term potentiation (LTP), and descending facilitation are among the best characterized forms of neuronal malplasticity proposed for OIH expression ^6, 9, 10, 60–62^. These and other lines of evidence suggest the importance of malplasticity of excitatory systems in OIH expression^9^. However, because functional homeostasis of normal neural circuits is maintained by the coordinated activity of excitatory and inhibitory neurons, it is also important to understand if and how this coordination is altered during OIH development. Yet, we currently know little in this aspect. We find that morphine indeed induces hyper-activity of excitatory inputs on excitatory neurons and hypo-activity of excitatory inputs on inhibitory neurons (Fig. 2). The increased synaptic drive on excitatory neurons and decreased synaptic drive on inhibitory neurons reveal a form of morphine-induced polarization in the SDH neural circuit. This form of neural circuit polarization (NCP) would facilitate hyper-activation of the pain neural circuit in the SDH and thus promote the expression of OIH. This neural circuitry mechanism may also play a critical role in HIV-1 gp120-induced pain pathogenesis^63^.

The increase of excitatory synaptic drive on excitatory neurons is consistent with the involvement of sensitization of pain processing neurons in the SDH during OIH expression, as suggested by previous studies^9^. It is unexpected for us to observe the decrease of excitatory synaptic drive on inhibitory neurons. Inhibitory loss, manifested as loss of inhibitory neurons or synapses, has been suggested previously as a potential mechanism underlying pain pathogenesis^64–66^. Our findings reveal novel form of loss of inhibition implicated in OIH development.

Our finding of NCP reveals a novel circuit mechanism that supports the expression of OIH. The increase excitatory inputs to excitatory neurons and the decrease of excitatory inputs to inhibitory neurons in NCP may promote OIH by facilitating hyper-activation of pain transmission circuits.

### Astrocytic mechanism in regulation of OIH expression

Previous studies on OIH mechanism mainly concentrate on the contribution of neuronal plasticity. The involvement of glial cells is recognized later, as indicated by the activation of microglia and astrocytes in rodent OIH models^9^. Microglia were suggested to modulate neuronal activity via BDNF during OIH expression^11^. However, other studies raise questions about the role of microglia^4, 12^. Astrogliosis is associated with various chronic pain conditions, including OIH models, but its pathogenic contribution has not been tested conclusively ^9, 16^. Previous work suggests a contribution of activated astrocytes to pain development in animal models^67, 68^. However, a recent study by Sasaki et al. show that inhibition of astrocyte activation by gap junction blockade did not ameliorate OIH^69^. It is important to note that the GFAP upregulation induced by morphine was not blocked in this study, raising uncertainty about the degree of astrogliosis inhibition.

The current speculations of astrocytic function in pain pathogenesis are still largely based on circumventive evidence such as the correlation between astrogliosis and pain and the effect of glial inhibitors^67–69^. We take a genetic approach to specific ablate reactive astrocytes and thus directly evaluate the contribution of reactive astrocytes to OIH. We show that astrogliosis ablation during the early phase of OIH induction impaired the expression of mechanical OIH (Fig. 1C), whereas astrogliosis ablation after the establishment of OIH significantly reversed the OIH (Fig. 1D). These results strongly suggest an important role of reactive astrocytes in both expression and maintenance of OIH.

Our conclusion described above is also supported by the effect of astrogliosis inhibition induced by the CKO of Wnt5a or ROR2. We show that when morphine-induced astrogliosis is blocked by either neuronal Wnt5a CKO or astrocytic ROR2 CKO OIH is abolished (Fig. 3; Fig. 4). These cell-type-specific approaches of astrogliosis inhibition provide strong complementary evidence for a crucial contribution of reactive astrocytes to OIH.

How would reactive astrocytes promote OIH? Little is known about the answer to this important question. Astrocytic processes intimately interact with synapses to form a tripartite synaptic structure^15^. Under physiological conditions, normal astrocytes play key roles in neural circuit homeostasis, by maintaining cation (e.g. K+) equilibrium, uptaking released neurotransmitters and releasing gliotransmitters^70^. However, it is unclear how reactive astrocytes would disturb the circuit homeostasis, especially in during OIH development. We find that morphine induces NCP and ablation of astrogliosis suppresses the NCP expression (Fig. 2). These findings suggest that morphine-induced reactive astrocytes disturb circuit homeostasis and mediate the expression of NCP. This notion is supported by the findings that inhibition of astrogliosis in either the neuronal Wnt5a CKO or the astrocytic ROR2 CKO mouse also blocks NCP expression (Fig. 3; Fig. 4). Based on these results, we propose that reactive astrocytes facilitate OIH by disrupting homeostasis of neural circuits in the SDH and promoting NCP expression.

Then, how would reactive astrocytes promote NCP? Based on the characterized biological activity of reactive astrocytes, it has been speculated that they could facilitate OIH expression by releasing excitatory substances to stimulate synapses. The suggested excitatory factors include proinflammatory cytokines and chemokines, ATP and gliotransmitters^9^. However, this hypothesis has not been directly and systematically tested in the context of OIH. Our results suggest that the reactive astrocytes in the OIH models generate active IL-1β (Fig. 5). We further show that blockade of IL-1 signaling by IL-1Ra impairs both OIH and NCP (Fig. 6). Based on these findings, we suggest that reactive astrocytes contribute to NCP via IL-1β.

### Wnt signaling in regulation of astrogliosis during OIH pathogenesis

Mechanism of astrocyte activation during OIH pathogenesis remains poorly understood. We find that neuronal Wnt5a and astrocytic ROR2 receptors are critical for morphine to activate astrocytes in the spinal cord (Fig. 3; Fig. 4). These findings suggest that Wnt5a secreted from neurons stimulates its receptor ROR2 on astrocytes to activate astrocytes. Our results also show that this neuron-to-astrocyte Wnt5a-ROR2 signaling pathway is required for OIH and NCP (Fig. 3; Fig. 4), both of which are dependent on reactive astrocytes (Fig.1; Fig. 2).

Wnt5a expression and secretion are evoked by synaptic activity^45^ and pain^44^. In addition, morphine is known to upregulate Wn5a in the spinal cord^47^. Hence, we propose that morphine treatment causes Wnt5a release from neurons to activate astrocytes. The requirement of neuronal Wnt5a indicates that morphine is not sufficient to activate astrogliosis in vivo, during OIH expression. These findings are consistent with the observation that μ-opioid receptor (MOR), which is critical for OIH development and critical for astrocyte activation induced by ultra-low-dose morphine ^14^, was reported not to be expressed in spinal astrocytes ^71^. However, because opioid receptors were detected in cultured astrocytes^72^ and morphine modulated astrocytic physiology in vitro^73, 74^, it is possible that morphine stimulates resting astrocytes but the stimulation is not sufficient to elicit full astrogliosis, which requires the facilitation of Wnt5a from neurons. It will be interesting to investigate this possibility in future studies. In this context, it is interesting to note that another Wnt family member Wnt3a is also involved in OIH, by modulating fractalkine/CX3CR1 in rats^75^. Hence, different Wnt signaling pathways may contribute to OIH development by modulating astrocytes and microglia.

Our data further show that Wnt5a receptor ROR2 on astrocytes is critical for morphine to activate astrocytes (Fig. 4). JNK signaling is an important intracellular mediator of the Wnt5a-ROR2 pathway^76^ and the JNK signaling are critical for astrocyte activation induced in OIH and neuropathic pain models ^14, 49^. Our data show astrocytic ROR2 CKO blocked morphine-induced phosphorylation of JNK in spinal cord (Fig. 7E) and JNK inhibitor also suppressed induced increase of IL-1 and casp-1 (Fig. 7F) These findings suggest that Wnt5a-ROR2 signaling mediated morphine-induced inflammasome activation via JNK signaling.

The neuron-to-astrocyte Wnt5a-ROR2 intercellular signaling pathway is also critical for spinal astrocyte activation induced by gp120 in an HIV pain model^63^, indicating that this pathway is a general mechanism that controls astrocyte activation induced by different neurological conditions.

Collectively, our results suggest that astrogliosis is a key cell target for OIH intervention. Furthermore, the results also suggest that the Wnt5a-ROR2 signaling pathway is a potential molecular target to block the astrogliosis during OIH pathogenesis. Ample clinical and preclinical evidence suggests the close relationship between OIH and tolerance. It will be interesting to test if the glial and molecular OIH mechanisms that we elucidate here also play critical role in the development of opioid tolerance.

## ACKNOWLEDGEMENT

We are grateful for the productive discussion and insights from Dr. Ye Zhang. This work was supported by NIH grants R01NS079166 (SJT), R01DA036165 (SJT), R01NS095747 (SJT), 1R01DA050530 (SJT, JMC) and 1R01NS122571 (SJT).

## AUTHOR CONTRIBUTIONS

Experimental design: SJT; Data collection and analysis: XL, CB, BL, YZ; Manuscript preparation: XL, SJT, XZ, JMC, TY

## SUPPLEMENTAL FIGURES AND LEGENDS

**Supplemental Figure 1.**
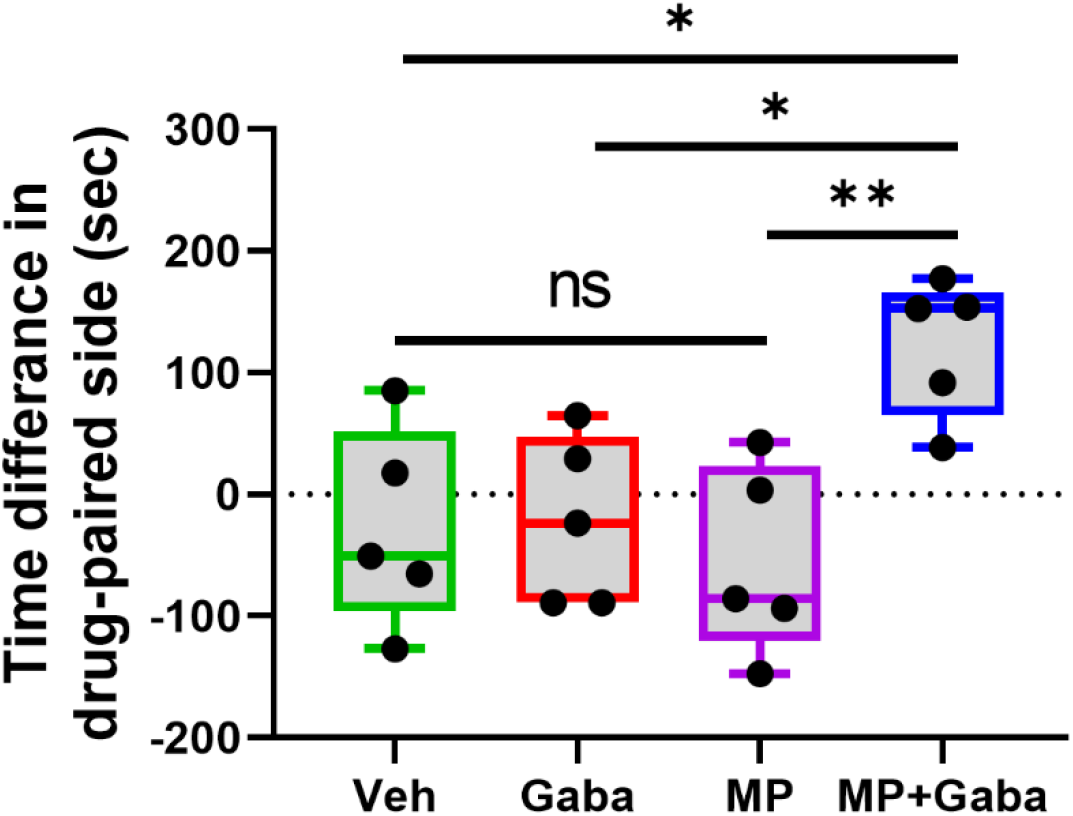
Mice repeatedly exposed to morphine developed a conditioned place preference for gabapentin. Data show that gabapentin (100mg/kg) injection developed conditioned place preference only in morphine-exposed mice compared with morphine-exposed alone (P<0.01), gabapentin-or vehicle treated alone mice (P<0.05) (n=5/group).

**Supplemental Figure 2.**
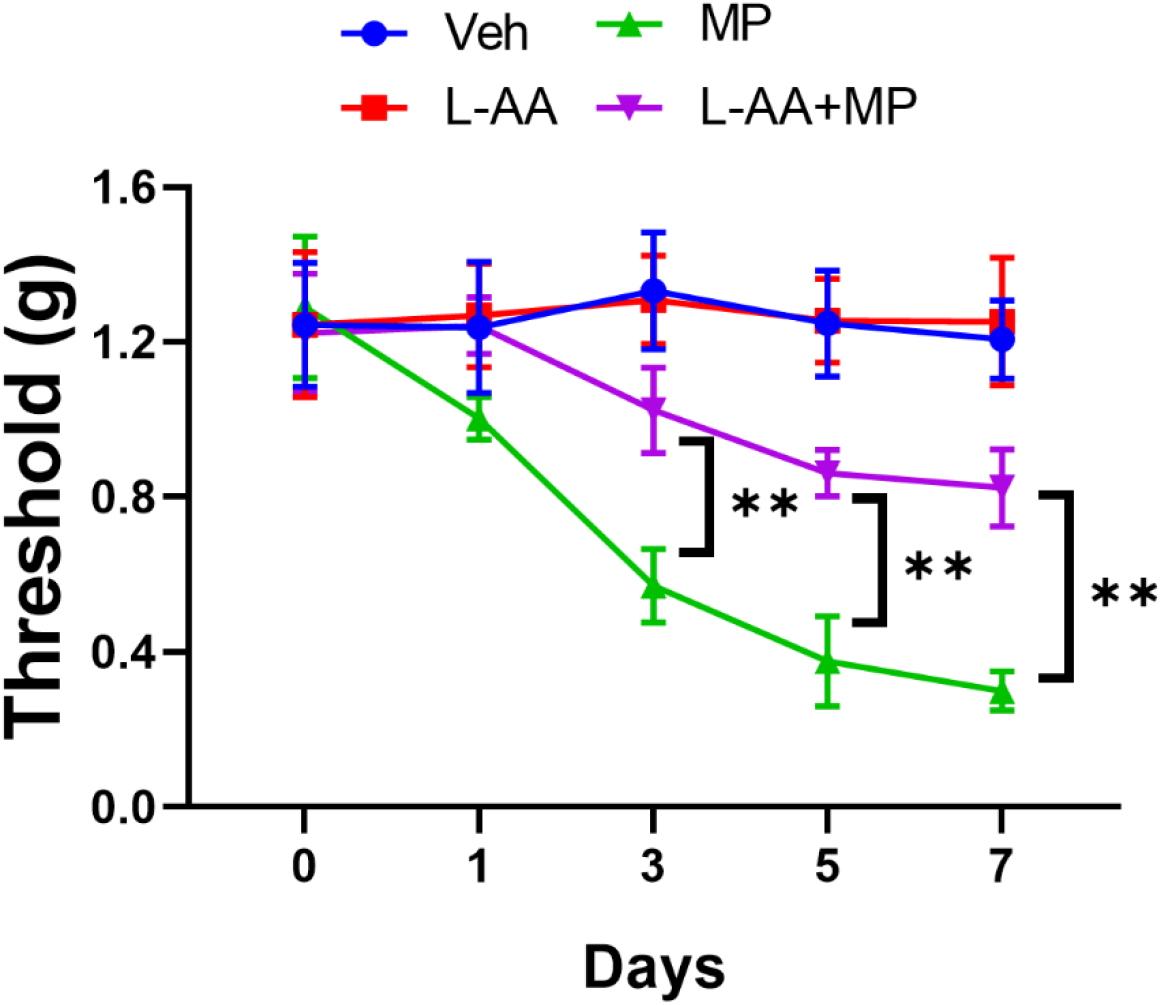
Effect of astrogliosis pharmacological ablation on OIH in female mice. Female wild type mice expressed mechanical OIH. Injections of L-AA (100nmol/5μl; i.t.) inhibited the morphine-induced increase of mechanical sensitivity, measured by von Frey tests (n=6/group).

**Supplemental Figure 3.**
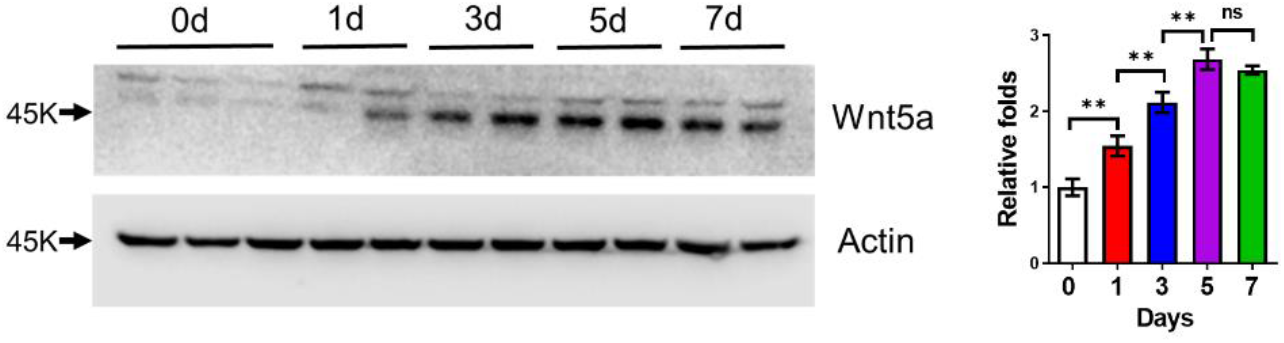
Immunoblotting analysis of spinal Wnt5a in wild type mice treated with morphine at different time course. Morphine was administered (20mg/kg) i.p. once daily. Mice were euthanized for spinal cord collection at day 0, 1, 3, 5 and 7 (n=3/group).

**Supplemental Figure 4.**
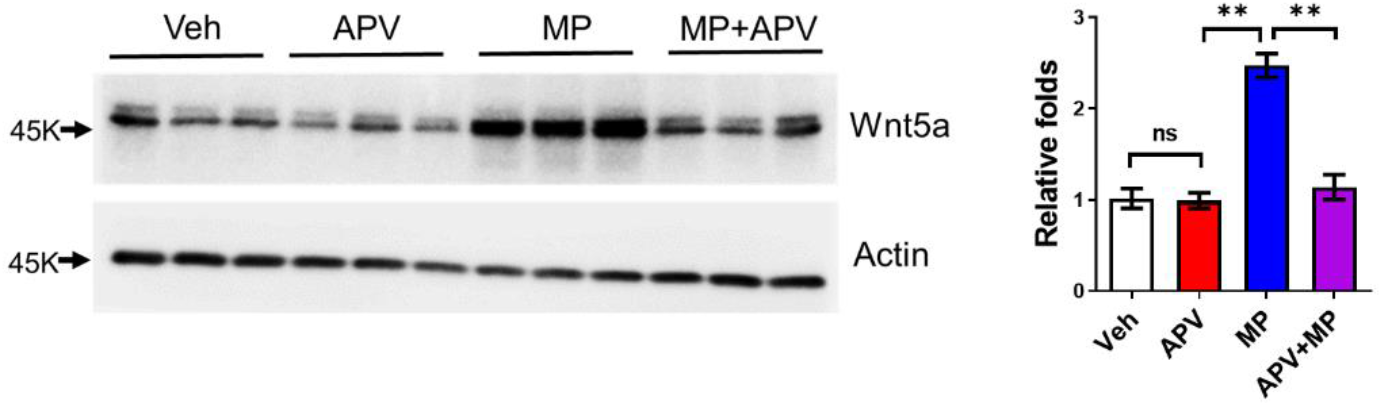
Synaptic activity is required for morphine induces spinal Wnt5a increase. Immunoblotting analysis of Wnt5a in the spinal cord of wild type mice treated with saline, APV, morphine or APV+morphine. NMDAR antagonist APV was intrathecally administered (5μg/5μl) at 0,2,4,6 day and 30 minutes prior to morphine injection. Mice were euthanized for spinal cord collection at day 7 (n=4/group).

**Supplemental Figure 5.**
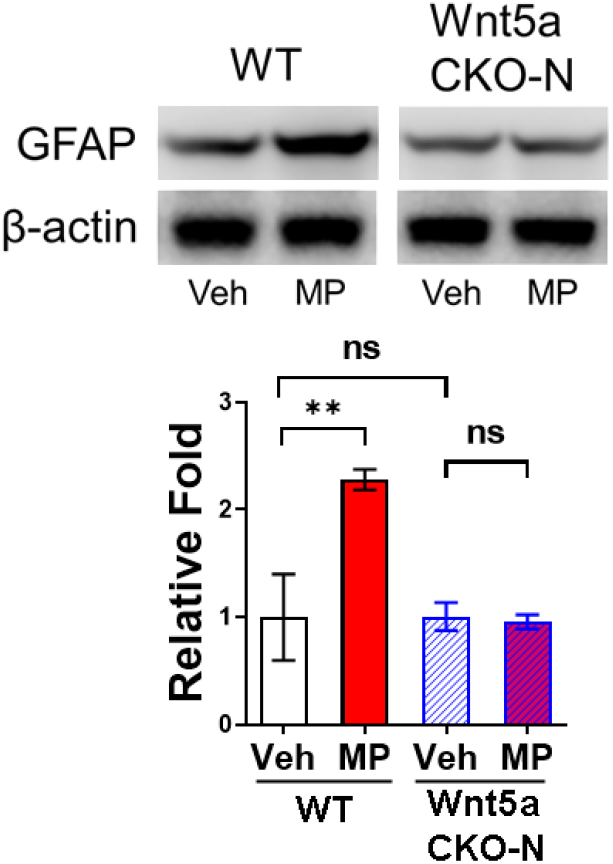
Wnt5a CKO-N blocked morphine-induced astrogliosis in the spinal cord.

**Supplemental Figure 6.**
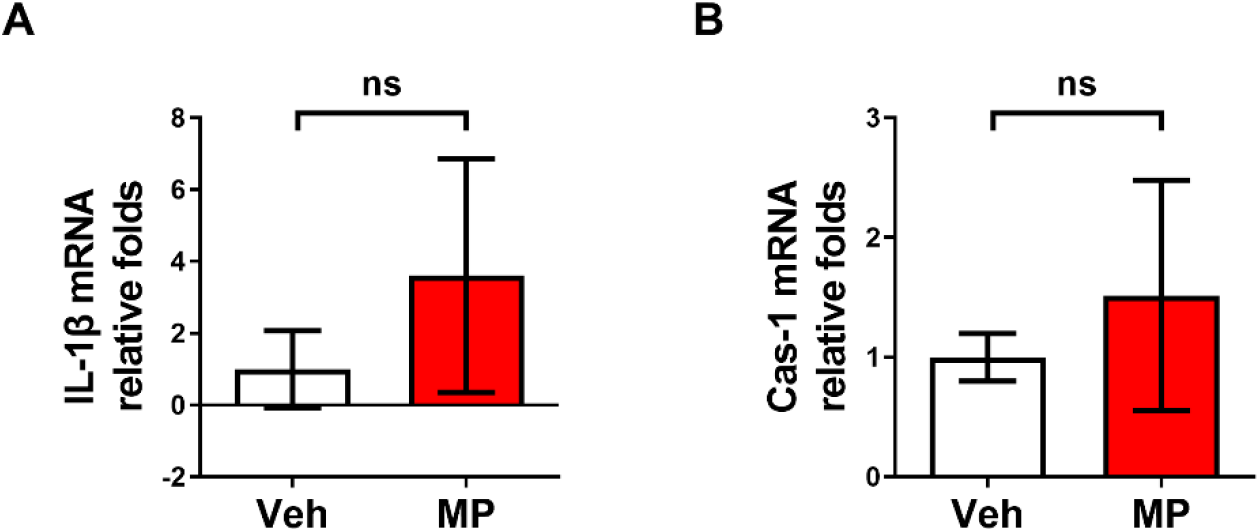
Morphine exposure did not induce the production of IL-1β and Cas-1 mRNA. Morphine administration paradigm showed in Fig. 1C. Mice were euthanized for spinal cord collection at day 7(n=5/group).

**Supplemental Figure 7.**
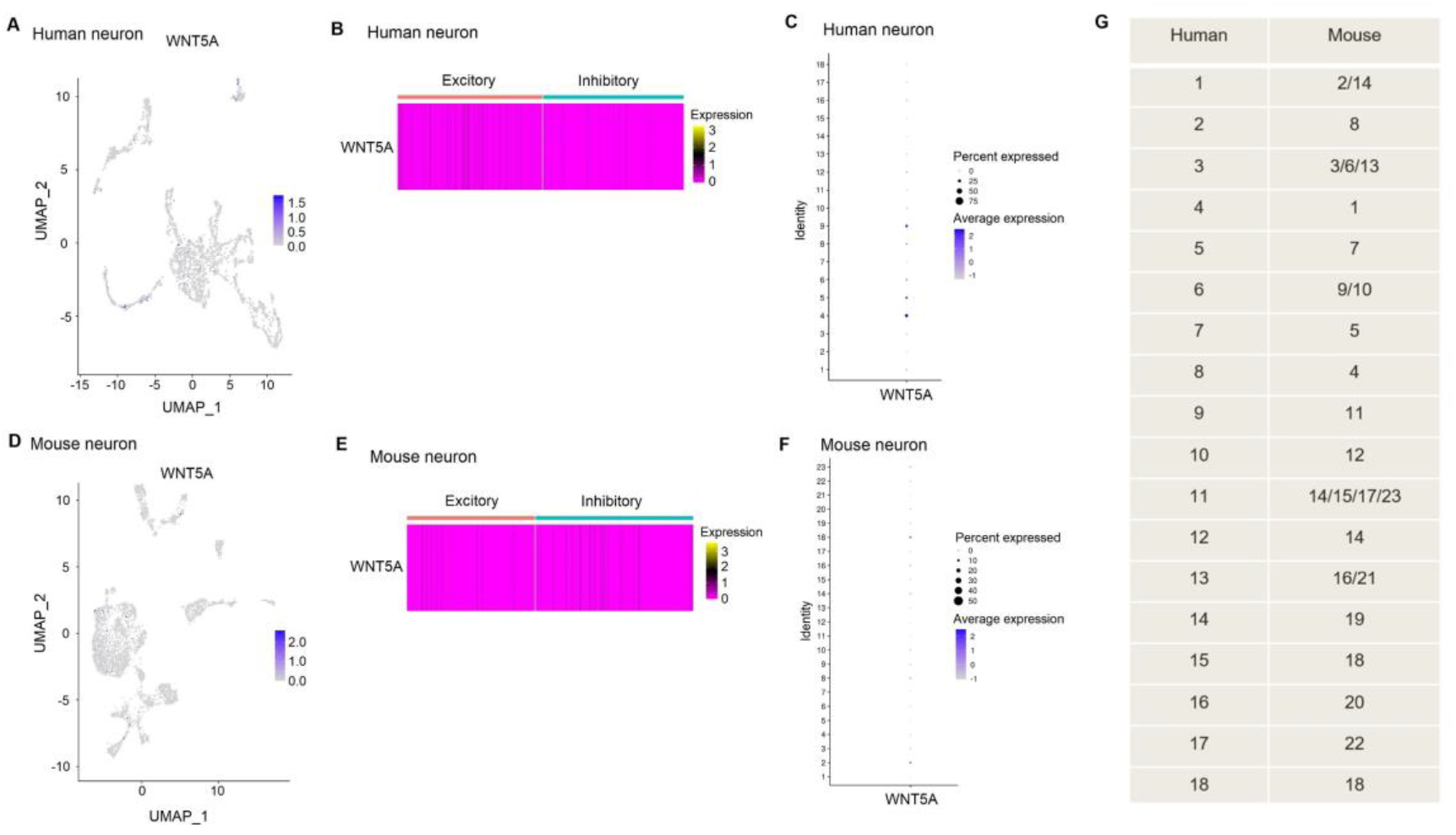
Expressional profiles of WNT5A in spinal neurons between humans and mice. A-C: UMAP plot (A), heatmap (B), and dot plot (C) showing the expressions of WNT5A across all neuronal types in human spinal cord. D-F: UMAP plot (D), heatmap (E), and dot plot (F) showing the expressions of WNT5A across all neuronal types in mouse spinal cord. G: The putative homologous neuronal clusters between human and mouse. UMAP, Uniform Manifold Approximation and Projection.

**Supplemental Figure 8.**
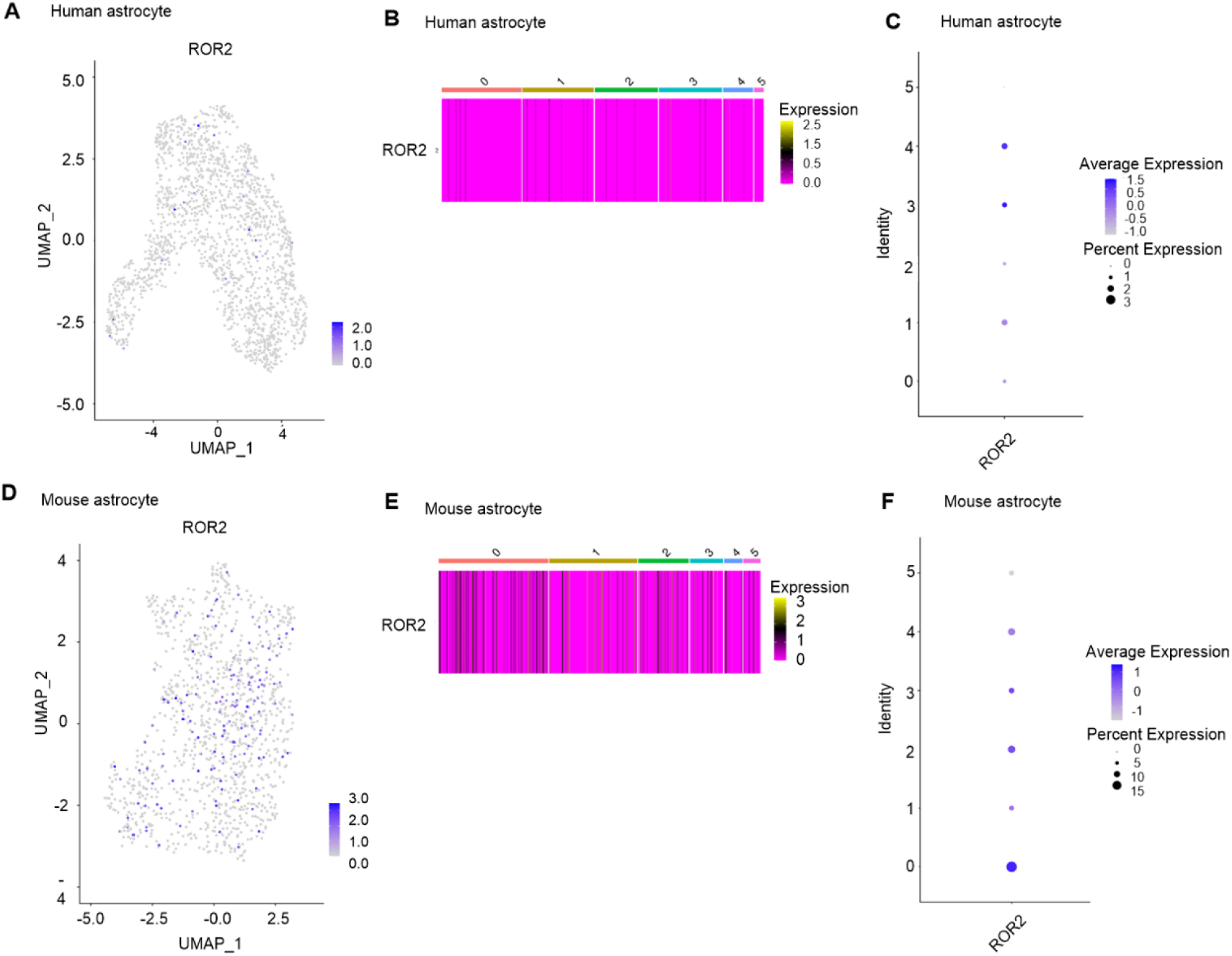
Expressional profiles of ROR2 in spinal astrocytes between humans and mice. A-C: UMAP plot (A), heatmap (B), and dot plot (C) showing the expressions of ROR2 across all astrocytic subtypes in human spinal cord. D-F: UMAP plot (D), heatmap (E), and dot plot (F) showing the expressions of ROR2 across all astrocyte subtypes in mouse spinal cord. UMAP, Uniform Manifold Approximation and Projection.

